# Blocking apoptosis promotes survival and alters developmental dynamics of human retinal ganglion cells in retinal organoids

**DOI:** 10.1101/2025.09.27.678991

**Authors:** Jingliang Simon Zhang, Brian Guy, Clayton P. Santiago, Caterina Tiozzo, Meghana Sreenath, Ya-Wen Chen, Seth Blackshaw, Robert J. Johnston

## Abstract

Retinal ganglion cells (RGCs) are the projection neurons that transmit visual information from the retina to the brain. In many species, a substantial proportion of RGCs are eliminated by programmed cell death during development to regulate their final number, but how cell death impacts human RGC development remains poorly understood. Here, we characterized the timing and cell-type-specificity of cell death in human fetal retinas and retinal organoids. Both retinas and organoids exhibited two waves of apoptosis: an early wave targeting neurogenic retinal progenitor cells and neuronal precursors, and a late wave affecting RGCs and other neurons. Additionally, organoids displayed a distinct wave of necrosis. To investigate how the apoptotic waves affect retinal development, we differentiated human BAX/BAK double mutant organoids deficient in apoptosis. In these mutants, RGC lifespan and survival increased, while RGC neurogenesis and maturation were delayed. Thus, developmental apoptosis controls not only the quantity of RGCs but also their developmental dynamics. Together, our results highlight the roles of apoptosis in human RGC development and the challenges in retinal organoid design. Addressing these limitations will improve the utility of organoids for studying human retinal development and modeling optic neuropathies like glaucoma.

## Introduction

The human retina is a laminated structure in the eye that enables vision. In the adult human retina, cell bodies of retinal neurons are distributed in three nuclear layers, including the outer nuclear layer (ONL), the inner nuclear layer (INL) and the ganglion cell layer (GCL) (Quinn and Wijnholds, 2019; Hussey et al., 2022). Located in the GCL, RGCs are the output neurons of the retina, projecting axons through the optic nerve to transmit visual information to the brain (Laha et al., 2017). As the first neuronal cell type specified in the human retina (Nguyen-Ba-Charvet and Rebsam, 2020), RGCs are overproduced during early development (Provis et al., 1985b). Whereas the majority of RGCs die later in development, a subset of RGCs survive, establishing their topography across the retina and stable retinotopy to the brain (Provis and Vandriel, 1985; Provis, 1987). Here, we study how programmed cell death regulates human RGC neurogenesis, maturation, and survival.

Programmed cell death is a conserved mechanism that regulates cell number and refines neuronal connectivity in the retina and other parts of the central nervous system (Oppenheim, 1981; Buss et al., 2006; Francisco-Morcillo et al., 2014; Yamaguchi and Miura, 2015). Two distinct waves of programmed cell death occur in the developing retina of many species (Glucksmann, 1940; Young, 1984; Gaze and Grant, 1992; Frade et al., 1997; Hensey and Gautier, 1998; Pequignot et al., 2003; Candal et al., 2005). The first wave occurs in the neuroblastic layer (NBL) and coincides with RGC neurogenesis (Frade et al., 1996; Pequignot et al., 2003; Candal et al., 2005). The second wave occurs during RGC innervation and maturation, eliminating RGCs that fail to correctly innervate their targets (Hughes and McLoon, 1979; Perry and Cowey, 1982; Rakic and Riley, 1983; Carpenter et al., 1986; Vanselow et al., 1990). Inhibition of apoptosis in the developing retina results in reduced developmental cell death, thickened retinal layers, increased RGC numbers, and impaired retinal functions (Cecconi et al., 1998; Mayordomo et al., 2003; Pequignot et al., 2003; Hahn et al., 2003; Ke et al., 2018). Thus, developmental apoptosis regulates RGC population size to ensure retinal function.

Our understanding of how cell death occurs and regulates human retinal development is limited. Human RGC neurogenesis starts in fetal week 6 and likely concludes by week 14, based on the temporal expression of key RGC developmental genes such as *ATOH7* and *POU4F2* (Hoshino et al., 2017; Aldiri et al., 2017). Caspase 3-expressing cells are observed in human retinas between weeks 5 and 8, suggesting an early wave of apoptosis (Bozanic et al., 2003). A second apoptotic wave occurs later, with a peak between gestation weeks 16 and 20 (post-conception weeks 14-18), when substantial loss of optic nerve axons and pyknosis of cells in the GCL are observed (Provis et al., 1985b; Provis et al., 1985a; Provis, 1987; Georges et al., 1999). This coincides with the stage when RGCs are sending axons to innervate their targets in the brain (Gilbert, 1935; Cooper, 1945; Khan et al., 1994; Hevner, 2000), suggesting that this second wave of cell death may refine the eye-to-brain connectivity by eliminating excess RGCs. However, the precise identity of the apoptotic cells and the specific roles of cell death in human RGC development have not been studied.

To address these questions, we studied cell death during the development of human fetal retinas and retinal organoids. Human stem cell-derived retinal organoids form optic vesicle-like structures (OVs) via evagination, generate all major neuronal cell types in the retina, and recapitulate early human retinal development *in vitro*, providing a highly accessible and experimentally tractable system for developmental and translational studies (Meyer et al., 2011; Nakano et al., 2012; Zhong et al., 2014; Cowan et al., 2020). In organoids, RGCs are generated in the inner layers and display neurite outgrowth, mimicking early RGC development *in vivo* (Fligor et al., 2018; Capowski et al., 2019). However, RGCs in organoids are almost completely lost during long-term culture, compromising studies of RGC biology and their potential for translational applications (**Fig. 1A**) (Capowski et al., 2019; Wahle et al., 2023). It has been suggested that RGCs are lost in organoids due to the lack of innervation targets, neurotrophic support, vasculature, and/or inaccessibility of oxygen in the innermost region (Fligor et al., 2021; Gong et al., 2023; Drabbe et al., 2025; Inagaki et al., 2025). Nevertheless, what drives RGC loss and how inhibition of apoptosis affects RGC development and maintenance in human organoids have not been examined.

**Figure 1.**
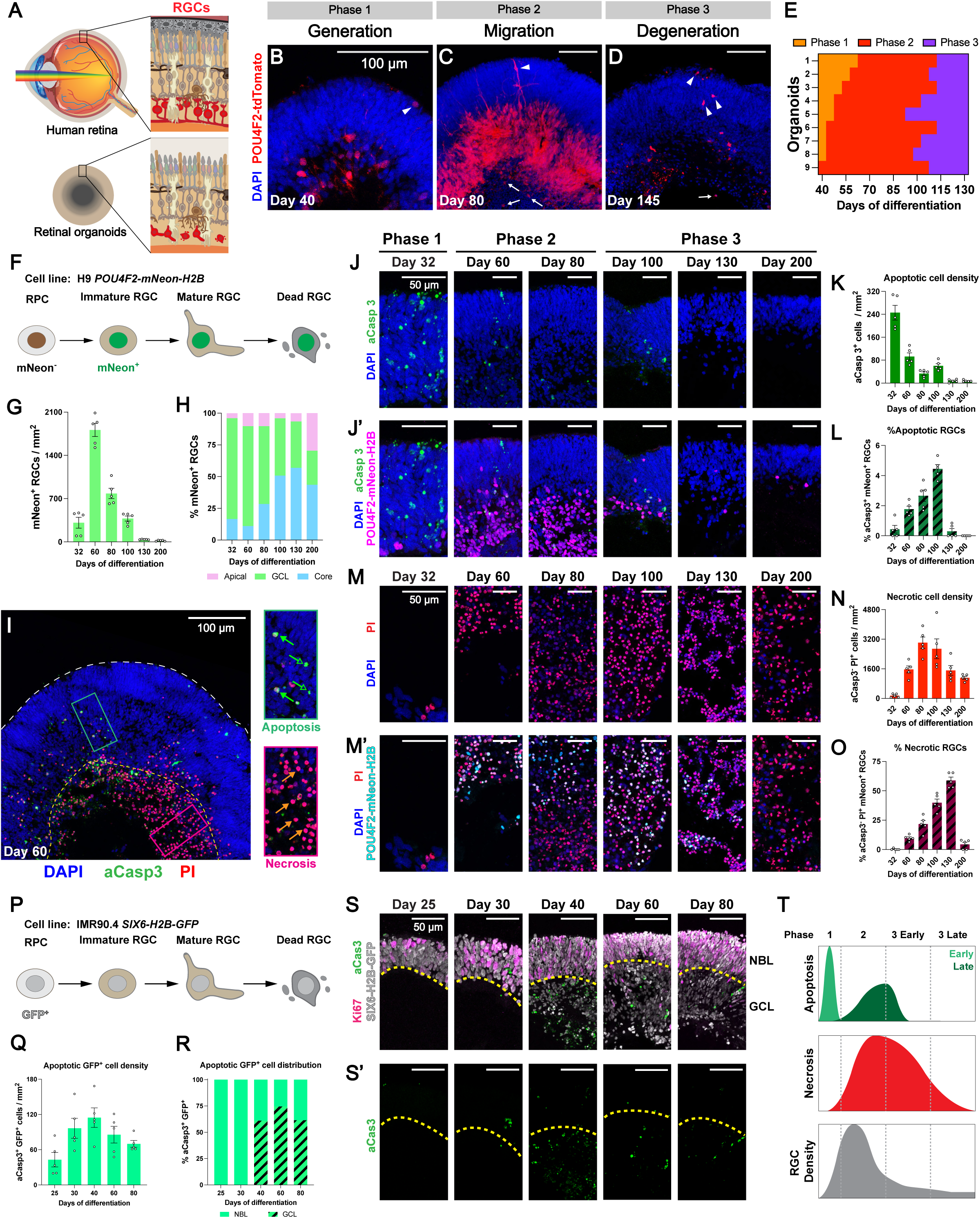
Waves of cell death in human retinal oranoids. **(A)** Schematic of RGCs (red) in the human retina and their loss in human retinal organoids. **(B-D)** Human retinal organoids in Phase 1 (B), Phase 2 (C), and Phase 3 (D) with distinct abundances and distributions of tdTomato^+^ RGCs (red). White arrowheads indicate mislocalized RGCs in the apical layers. White arrows indicate mislocalized RGCs in the core. **(E)** Progression of the three phases of RGC development in human retinal organoids based on live imaging. **(F)** As *POU4F2* is expressed in post-mitotic RGC precursors, *POU4F2-p2A-mNeon-H2B* labels immature, mature, and dead RGCs, but not RPCs. **(G)** Density of mNeon^+^ RGCs in human retinal organoids. Data are presented as mean ± SEM. N = 5 organoids per timepoint. **(H)** Distribution of mNeon^+^ RGCs in different layers or regions of human retinal organoids. N = 5 organoids per timepoint. **(I)** Co-detection of apoptosis and necrosis in H9 *POU4F2-mNeon-H2B* retinal organoids. Empty green arrows indicate aCasp3^+^ PI^-^ cells (early apoptotic cells) and solid green arrows indicate aCasp3^+^ PI^+^ cells (late apoptotic cells) in retinal layers. Orange arrows indicate aCasp3^+^ PI^-^ cells (necrotic cells) in the core. **(J)** Apoptotic cells (aCasp3^+^) and apoptotic RGCs (aCasp3^+^ mNeon^+^) in retinal layers. **(K-L)** Density of aCasp3^+^ cells in retinal layers (K) and % aCasp3^+^ mNeon^+^ RGCs of all mNeon^+^ RGCs (L). Data are presented as mean ± SEM. N = 5 organoids per timepoint. **(M)** Necrotic cells (PI^+^) and necrotic RGCs (PI^+^ mNeon^+^) in the core. **(N-O)** Density of aCasp3^-^ PI^+^ cells in the core (N) and % aCasp3^-^ PI^+^ mNeon^+^ RGCs of all mNeon^+^ RGCs (O). Data are presented as mean ± SEM. N = 5 organoids per timepoint. **(P)** *SIX6-p2A-H2B-GFP* labels RPCs, as well as immature, mature, and dead RGCs. **(Q-R)** Density of aCasp3^+^ GFP^+^ cells in retinal layers (Q) and distribution of aCasp3^+^ GFP^+^ cells in the NBL and GCL (R) during the early apoptotic wave. For Q, data are presented as mean ± SEM. N = 4 organoids per timepoint. **(S)** Detection of retinal cells (GFP^+^) and apoptotic retinal cells (aCasp3^+^ GFP^+^) during the early wave of apoptosis. The yellow dashed line indicates the boundary of the NBL and GCL based on Ki67 (magenta) expression in the NBL. **(T)** Schematic of RGC density dynamics and waves of apoptosis and necrosis during human retinal organoid differentiation.

Here, we investigated the spatiotemporal dynamics and functional roles of cell death in human fetal retinas and retinal organoids using immunofluorescence microscopy, live imaging, birthdating, and single-nucleus RNA sequencing (scRNA-seq). Our study defines the temporal waves and regulatory roles of cell death during human retinal development, providing a foundation for understanding the intricate mechanisms that shape retinal cell populations and advancing organoid model design to study neuro-degenerative diseases.

## Results

### Two waves of developmental apoptosis in the human fetal retina

Considering the challenges in acquiring and analyzing fetal human retina samples, previous analyses of retinal apoptosis were limited to time windows that only encompassed a single wave of developmental apoptosis, and they relied on morphological features rather than cell type-specific markers to determine the identity of apoptotic cells (Provis et al., 1985b; Provis, 1987; Bozanic et al., 2003). To validate the timing of the two waves of apoptosis and the identities of apoptotic cells during human retinal development, we evaluated postmortem human fetal retinas collected at post-conception weeks (PCW) 9, 13, 16, and 20 in parasagittal sections (**Fig. S1A-D**). Apoptotic cells were detected by immunofluorescent staining for active Caspase 3 (aCasp3), the cleaved form of Caspase 3 generated during apoptosis (**Fig. S1A’-D’**) (Porter and Janicke, 1999). To identify apoptotic RGCs from other apoptotic cells in the GCL (*e.g.*, starburst amacrine cells) (Rodieck and Marshak, 1992), we co-stained with RBPMS, an RGC-specific marker in the mammalian retina (**Fig. S1A-D**) (Rodriguez et al., 2014; Yan et al., 2020).

In the NBL, aCasp3^+^ cells were primarily observed at PCW 9 (1.5 cells/mm) and decreased by PCW 13 (0.2 cell/mm) (**Fig. S1A-B, S1E-F**), reminiscent of the first wave of developmental apoptosis affecting RPCs and neuronal precursors in other vertebrate species (Frade et al., 1996; Frade, 2000; Pequignot et al., 2003).

In the GCL, the density of apoptotic cells was low at PCW 9 (0.4 cell/mm), increased by PCW 16 (2.5 cells/mm), and then decreased by PCW 20 (0.4 cell/mm) (**Fig. S1E-F**), marking the second wave of developmental apoptosis in human fetal retina. Importantly, we observed apoptotic cells in the GCL that expressed RBPMS (18 out of 3900 RBPMS^+^ RGCs examined at PCW 13, 42 out of 4365 RBPMS^+^ RGCs examined at PCW 16) (**Fig. S1B’-C’, S1G-H**), providing direct evidence of RGC apoptosis during human retinal development.

To evaluate the spatial dynamics of developmental apoptosis in the human fetal retinas, we quantified the density of apoptotic cells and their distribution along the vertical meridian (**Fig. S1I-K**). We found a high density of apoptotic cells in the NBL at PCW 9, and it decreased by PCW 13 as the fetal retina grew in length (**Fig. S1I**). We also observed a shift in the peak density of apoptotic RGCs from the central to the peripheral retina between PCW 13 and 16 (**Fig. S1J**), followed by the establishment of a central peak of RGC density at PCW 20 (**Fig. S1K**), consistent with analysis of horizontal sections of developing human retinas (Provis et al., 1983; Provis, 1987). In addition, we detected apoptotic cells in the INL, supporting previous reports of INL cell death during the second wave of apoptosis (**Fig. S1C’, S1F**) (Cook et al., 1998; Georges et al., 1999; Pequignot et al., 2003; Chavarria et al., 2007).

Together, these findings suggest that two waves of developmental apoptosis, an early wave affecting cells in the NBL and a later wave involving RGCs and other neurons, contribute to the sculpting of neuronal topography during human retinal development.

### Three phases of RGC development and loss in human retinal organoids

Human retinal organoids provide a highly accessible and experimentally tractable model to study human RGC development (Langer et al., 2018; Fligor et al., 2018; Fligor et al., 2021). To track changes in the spatial distributions and morphologies of RGCs during organoid development, we differentiated human retinal organoids from embryonic stem cells carrying the RGC-specific transgenic reporter POU4F2-p2A-tdTomato (H7 *POU4F2-tdTomato*) (Sluch et al., 2017) (**Fig. S2**). After 25 days of differentiation, a subset of retinal organoids formed OVs, the evaginating immature retinal tissue (**Fig. S3A**). By day 40, OVs were excised from the non-retinal parts and maintained in culture until day 200, while organoids lacking OVs were culled. Most OVs formed laminated retinal layers, including the GCL on the basal side, other retinal layers on the apical side (NBL between days 25-100 or ONL and INL between days 100-200), and a core with low cell density in the innermost region during culture (**Fig. S3B**).

Our observations revealed three phases of RGC development and loss in retinal organoids (**Fig. 1B-D**): generation (Phase 1), migration (Phase 2), and degeneration (Phase 3). These phases are distinct from the three general stages of human retinal organoid development previously characterized based on organoid morphology (Capowski et al., 2019).

In Phase 1 organoids, most tdTomato^+^ RGCs were in the putative GCL at the basal side of the retinal layers (**Fig. 1B**). In Phase 2 organoids, tdTomato^+^ RGCs started extending neurites. During this phase, although most RGCs still remained in the GCL, a subset of RGCs were mislocalized along extending neurites in the apical NBL or in the core (**Fig. 1C**), suggesting migration from the GCL. Also, pyknotic nuclei accumulated in the core during Phase 2 (**Fig. 1C**). In Phase 3 organoids, fewer tdTomato^+^ RGCs were observed. RGCs appeared to retract their neurites and retain fluorescent signal mainly in their soma (**Fig. 1D**), indicative of degeneration. In Phase 2 and 3 organoids, we observed the progressive breaching and deformation of a laminin-enriched structure under the GCL resembling the internal limiting membrane (ILM) (**Fig. S3C-E**), the innermost retinal layer essential for RGC polarity and neurite growth (Zhang and Johnson, 2021). These results indicate a correlation between ILM disruption and RGC mislocalization to the core. The progressive loss of tdTomato^+^ RGCs in human retinal organoids aligns with previous studies using different methods of organoid differentiation (Capowski et al., 2019; Fligor et al., 2021), suggesting that RGC loss is a common feature of long-term retinal organoid culture.

To determine when retinal organoids progress through each phase of RGC development, we tracked the development of tdTomato^+^ RGCs in nine individual retinal organoids from day 40 to day 130 (**Fig. 1E, S3F**). While organoids began the observation period in different phases and the absolute timing of each phase varied, the relative timing and progression were consistent across all organoids (**Fig. 1E**). Phase 1 was relatively short, with a maximum duration of 25 days from the start of observation. Phase 2 was observed primarily between days 50 and 100. All organoids entered Phase 3 by day 110 and remained in this phase (**Fig. 1E**).

Taken together, these results demonstrate that RGCs in organoids develop from Phase 1 (generation, days 25-50) to Phase 2 (migration, days 50-100), and finally, to Phase 3 (degeneration, days 100-200) (**Fig. S3G-J**), providing a general timeline for RGC development and loss in human retinal organoids.

### Tracking RGCs with a permanent transgenic reporter in retinal organoids

The progressive loss of *POU4F2-p2A-tdTomato*-expressing human RGCs could be due to downregulation of *POU4F2* during normal RGC maturation (Luo et al., 2019; Kriukov et al., 2024) or cell death (Soto et al., 2008). To track RGCs throughout their lifespan, we differentiated human retinal organoids from embryonic stem cells carrying a stable transgenic RGC-specific nuclear reporter *POU4F2-p2A-mNeon-H2B* (H9 *POU4F2-mNeon-H2B*) (Agarwal et al., 2023). Because *POU4F2* is expressed in post-mitotic immature RGCs, and fluorescent proteins fused with histone proteins remain stably associated with chromatin in post-mitotic neurons (Kanda et al., 1998; Tayler et al., 2011; Wiese et al., 2013), this reporter permanently labels RGCs that expressed *POU4F2*, as they mature or undergo cell death (**Fig. 1F, S3K**). Moreover, the nuclear mNeon reporter facilitates accurate quantification of RGCs in organoids by minimizing confounding signals from overlapping RGC soma and neurites, which arise with the cytosolic tdTomato reporter (**Fig. 1C**).

To evaluate the dynamics of mNeon^+^ RGC abundance, we quantified the density of mNeon^+^ RGCs in organoids on days 32, 60, 80, 100, 145, and 200. The density was low on day 32 in Phase 1, peaked on day 60 in early Phase 2, and progressively declined from day 80 onwards in late Phase 2 and Phase 3 (**Fig. 1G**, all measurements of cell densities and proportions are also listed in **Supplementary Tables**). The loss of permanently labeled mNeon^+^ RGCs directly demonstrates RGC death and clearance during organoid culture, consistent with our previous observation in H7 *POU4F2-tdTomato* organoids (**Fig. 1B-D**).

To assess the dynamics of mNeon^+^ RGC migration, we quantified the percentage of mNeon^+^ RGCs in the GCL, apical layers (NBL or ONL+INL), and core. In Phase 1 on day 32, most mNeon^+^ RGCs were located in the GCL (**Fig. 1H**). In Phase 2 on day 60, the proportion of mNeon^+^ RGCs in the GCL decreased, while the proportions in the apical layers and core increased (**Fig. 1H**), consistent with RGC migration from the GCL (**Fig. 1C**). In Phase 3 from day 100 and on, the proportion of mNeon^+^ RGCs in the GCL continued to decline, while the proportion in the core increased between days 100 and 130 and then decreased by day 200 (**Fig. 1H**), suggesting distinct dynamics of RGC loss in the GCL and the core. Together, these results define the spatiotemporal dynamics of RGC neurogenesis, migration, and loss during human retinal organoid development.

### RGCs die by apoptosis and necrosis in retinal organoids

The loss of mNeon^+^ RGCs suggested that RGCs undergo cell death during long-term organoid culture. Apoptosis of RGCs occurs in developing human retinas (**Fig. S1B-C, S1H**) (Provis et al., 1985b; Provis et al., 1985a; Provis, 1987) and mouse stem-cell derived retinal organoids *in vitro* (Volkner et al., 2021). In addition, the migration and loss of mNeon^+^ RGCs in the core of OVs (**Fig. 1H, S3K**) is reminiscent of the accumulation of necrotic cells in the core of human brain organoid models (Lancaster et al., 2013; Berger et al., 2018; Qian et al., 2020). Thus, we hypothesized that RGC loss in human retinal organoids results from apoptosis and/or necrosis.

Propidium iodide (PI) is a cell-impermeable, red-fluorescent nuclear dye that marks the nuclei of cells undergoing membrane rupture during necrosis and late-stage apoptosis (Rieger et al., 2011; Crowley et al., 2016). Co-detection with apoptosis-specific markers and PI is commonly used in flow cytometry to distinguish apoptotic and necrotic cells (Rieger et al., 2011; Liu et al., 2012; Costigan et al., 2023), but this approach had not been applied to immunofluorescent staining of organoid sections. To distinguish between apoptosis and necrosis in human retinal organoids, we developed an imaging-based co-detection assay using the antibody against aCasp3 and the dye PI. While aCasp3 marks all apoptotic cells, PI labels nuclei of late-stage apoptotic cells and necrotic cells. Thus, we classified aCasp3^+^ PI^-^ cells as early-stage apoptotic cells, aCasp3^+^ PI^+^ cells as late-stage apoptotic cells, and aCasp3^-^ PI^+^ cells as necrotic cells (**Fig. 1I**). Detection of all three populations validated the ability of this co-detection assay to distinguish apoptotic and necrotic cells in organoid sections.

Terminal deoxynucleotidyl transferase dUTP nick end labeling (TUNEL) is commonly used to detect cell death in developing retinas (Hahn et al., 2003; Georges et al., 1999). We found that TUNEL marks both a fraction of apoptotic cells and necrotic cells in day 60 organoids, suggesting that TUNEL is not ideal for distinguishing between apoptosis and necrosis for this study (**Fig. S4A-B**).

We next examined the regional distribution of apoptotic and necrotic cells. In day 60 organoids, 65% of apoptotic cells were localized to the retinal layers, while 88% of necrotic cells were found in the innermost core of the OVs (**Fig. S4C**), suggesting that apoptosis primarily occurs in the retinal layers, while necrosis predominantly affects cells in the core. For subsequent analysis, we focused on apoptotic cells in the retinal layers and necrotic cells in the core.

To investigate the dynamics of cell death during organoid development, we quantified apoptotic and necrotic cells in OVs on days 32, 60, 80, 100, 130, and 200, spanning the three phases of RGC development and loss (**Fig. 1J-O**).

We first evaluated the overall densities of apoptotic cells (aCasp3^+^ PI^-^ and aCasp3^+^ PI^+^ cells) in the retinal layers during organoid development. Phase 1 organoids displayed the highest density of apoptotic cells (**Fig. 1J-K**, day 32), suggesting an early wave of apoptosis. The density of apoptotic cells decreased in Phase 2 (**Fig. 1J-K**, days 60 and 80) and increased again in early Phase 3 (**Fig. 1J-K**, day 100), forming a late wave of apoptosis. The density decreased later in Phase 3 (**Fig. 1J-K**, days 130 and 200). These early and late waves of apoptosis in retinal layers of OVs (**Fig. 1K**) resemble the two waves of developmental apoptosis in human (**Fig. S1**) and other vertebrate retinas (Glucksmann, 1940; Young, 1984; Gaze and Grant, 1992; Hensey and Gautier, 1998; Pequignot et al., 2003; Candal et al., 2005).

We then assessed the apoptosis of RGCs during organoid development by quantifying aCasp3^+^ mNeon^+^ RGCs. Phase 1 organoids displayed a low proportion of apoptotic mNeon^+^ RGCs (**Fig. 1J’, 1L**, day 32). The proportion and density of apoptotic mNeon^+^ RGCs increased in Phase 2 and early Phase 3 (**Fig. 1J’, 1L, S4D**, days 60, 80 and 100), followed by their decrease later in Phase 3 (**Fig. 1J’, 1L**, **S4D**, days 130 and 200). H7 *POU4F2-tdTomato* organoids displayed similar temporal patterns of apoptosis, showing reproducibility across cell lines (**Fig. S4E-G**). The highest proportion of apoptotic mNeon^+^ RGCs coincides with the increase in overall apoptosis on day 100 (**Fig. 1K-L**), suggesting that apoptosis of RGCs contributes to the late wave of apoptosis in organoids.

We next evaluated the overall densities of aCasp3^-^ PI^+^ necrotic cells in the core during organoid development. A small number of necrotic cells were observed in Phase 1 (**Fig. 1M-N**, day 32). The density of necrotic cells increased rapidly in Phase 2 and early Phase 3 (**Fig. 1M-N**, days 60, 80 and 100), then decreased later in Phase 3 (**Fig. 1M-N**, days 130 and 200). These results demonstrate one wave of necrosis in the core of OVs during organoid development.

We then assessed the necrosis of RGCs by quantifying aCasp3^-^ PI^+^ mNeon^+^ RGCs. Few necrotic mNeon^+^ RGCs were observed in Phase 1 (**Fig. 1M’, 1O**, day 32). The proportion and density of necrotic mNeon^+^ RGCs then increased in Phases 2 and Phase 3 (**Fig. 1M’, 1O**, **S4H**, days 60, 80, 100 and 130), followed by a sharp decrease in late Phase 3 (**Fig. 1M’, 1O, S4H**, day 200). These results suggest that mNeon^+^ RGCs undergo significant necrosis between Phases 2 and 3 in organoids.

To evaluate the contribution of apoptosis or necrosis to the loss of RGCs in retinal organoids, we compared the proportions of apoptotic and necrotic mNeon^+^ RGCs. Whereas <5% of mNeon^+^ RGCs were apoptotic (**Fig. 1L**), 10% to 59% of mNeon^+^ RGCs were necrotic between days 60 and 130 (**Fig. 1O**). Moreover, this period of high-level RGC necrosis coincides with the decline of mNeon^+^ RGC density (**Fig. 1G**), suggesting that necrosis is a predominant driver of RGC loss in human retinal organoids.

### Neurogenic RPCs and newborn neuronal precursors die during the early wave of apoptosis

Since RGC death occurs during the second wave of apoptosis and the wave of necrosis, other cell types likely die during the early wave of apoptosis. During the early wave of apoptosis in the chick retina, TUNEL^+^ cells were identified as proliferating neuroepithelial cells or migrating newborn RGC precursors (Frade et al., 1997; Frade et al., 1996; Diaz et al., 2000; Frade, 2000). Considering the similar localization of apoptotic cells in the NBL of human fetal retinas (**Fig. S1A’**) and organoids (**Fig. 1J**), we hypothesized that these apoptotic cells are RPCs or postmitotic neuronal precursors.

To label the cell types contributing to the early wave of apoptosis, we differentiated retinal organoids from a human induced pluripotent stem cell (iPSC) line carrying the transgenic nuclear reporter SIX6-p2A-H2B-GFP (IMR90.4 *SIX6-H2B-GFP*) (Wahlin et al., 2021). In organoid-derived OVs, H2B-GFP permanently labels all cells that expressed SIX6, including RPCs and their postmitotic neuronal progeny (**Fig. 1P**) (Toy and Sundin, 1999; Toy et al., 1998; Aijaz et al., 2005). To test if the SIX6-H2B-GFP reporter marks the apoptotic cells during the early wave, we measured the densities of aCasp3^+^ GFP^+^ cells in the retinal layers (*i.e.,* NBL and GCL) of IMR90.4 *SIX6-H2B-GFP* organoids on days 25, 30, 40, 60, and 80. We observed a peak of apoptotic aCasp3^+^ GFP^+^ cells on day 40 (**Fig. 1Q**), validating our previous observation that the early wave of apoptosis peaked between days 30 and 40 (**Fig. 1K**).

We next evaluated the distribution of apoptotic aCasp3^+^ GFP^+^ cells in the NBL and GCL during early organoid development. On days 25 and 30, nearly all aCasp3^+^ GFP^+^ cells were found in the NBL (**Fig. 1R-S**). During the formation of GCL, over 50% of aCasp3^+^ GFP^+^ were observed in the GCL on days 40, 60, and 80 (**Fig. 1R-S**). These data suggest that the early wave of apoptosis is composed of RPCs in the NBL first and immature neuronal precursors migrating to the developing GCL later.

To determine the contribution of RPCs to the early wave of apoptosis, we quantified the proportion of aCasp3^+^ GFP^+^ cells expressing the RPC marker VSX2 on days 30 and 40. Less than 10% of aCasp3^+^ GFP^+^ cells were VSX2^+^ on days 30 and 40 (**Fig. S4I-K**), suggesting that VSX2^+^ RPCs are not the major cell type undergoing apoptosis at this stage.

At the beginning of retinal neurogenesis, the NBL consists of RPCs—including proliferative RPCs (pRPCs) that highly express cell-cycle phase-associated genes and neurogenic RPCs (nRPCs) that upregulate proneural transcription factors for potential neurogenic division—as well as postmitotic neuronal precursors (Alexiades and Cepko, 1996; Clark et al., 2019; Shiau et al., 2021). To determine whether the apoptotic cells in the NBL are pRPCs, nRPC and/or neuronal precursors, we pulsed organoids with EdU (5-ethynyl-2’-deoxyuridine) for 24 hours on day 26 and measured the proportion of apoptotic aCasp3^+^ GFP^+^ cells that were Ki67^+^ (pRPCs) or EdU^+^ (recently divided pRPCs, nRPCs and newborn neuronal precursors) on day 30 (**Fig. S4I**). 41.1% of aCasp3^+^ GFP^+^ cells were EdU^+^ (**Fig. S4M-N**), while only 3.7% were Ki67^+^ (**Fig. S4M, S4O**), suggesting that the early wave of apoptosis in human retinal organoids is composed primarily of nRPCs and/or newborn neuronal precursors, with a smaller contribution from pRPCs.

In summary, we observed an early wave of apoptosis during Phase 1, a late wave of apoptosis in retinal layers during Phase 2 and early Phase 3, and a wave of necrosis in the core expanding Phases 2 and 3 in human retinal organoids. These results suggest that the early wave of apoptosis mainly affects nRPCs and neuronal precursors, while the late wave of apoptosis and necrosis lead to the substantial loss of RGCs during long-term culture (**Fig. 1T**).

### *BAX/BAK* knockout eliminates the two waves of apoptosis in retinal organoids

We next sought to investigate the roles of the two waves of apoptosis in human retinal organoids by inhibiting developmental apoptosis. BAX and BAK are pro-apoptotic proteins in the *BCL2* family that are essential regulators of the apoptotic pathway (Pena-Blanco and Garcia-Saez, 2018). In mouse retinas, genetically knocking out *Bax* and *Bak* inhibits developmental apoptosis, leading to increased thickness and cell numbers of the GCL and INL (Pequignot et al., 2003; Hahn et al., 2003). CRISPR-edited genetic knockout of *BAX* and *BAK* in human iPSCs inhibits apoptosis (**Fig. S5A**) (Joshi et al., 2020), providing an effective model to study the role of developmental apoptosis in human development.

To investigate how apoptosis affects retinal organoid development, we differentiated human retinal organoids from *BAX* and *BAK* double mutant (GM25256 *BAX/BAK dKO*) and isogenic control (GM25256) iPSCs (**Fig. S5B-C**) (Joshi et al., 2020). *BAX/BAK dKO* and control organoids displayed no significant differences in the sizes of OVs or their cores (**Fig. S5D-E**). Organization of the retinal layers was similar throughout differentiation, except at day 200, when the retinal layers in *BAX/BAK dKO* organoids were significantly thicker (**Fig. S5F**). These observations suggest that inhibition of apoptosis in *BAX/BAK dKO* organoids leads to improved long-term maintenance of retinal cells. Immunofluorescent staining confirmed the presence of major retinal neuronal types, indicating that *BAX/BAK dKO* does not block cell fate specification (**Fig. S5G-V**).

To assess how *BAX/BAK dKO* affects apoptosis in organoids, we quantified the densities of apoptotic cells (aCasp3^+^ PI^-^ and aCasp3^+^ PI^+^) (**Fig. 2A-B**). Control organoids displayed an early wave of apoptosis on days 40 and 60, and a late wave of apoptosis on day 100 (**Fig. 2A, 2C**), similar to our observations in organoids derived from other cell lines (**Fig. 1J-K, Fig. S4E**), while *BAX/BAK dKO* organoids displayed low densities of apoptotic cells at all timepoints (**Fig. 2B-C**). These results suggesting that *BAX/BAK dKO* essentially abolishes the two waves of apoptosis in retinal organoids.

**Figure 2.**
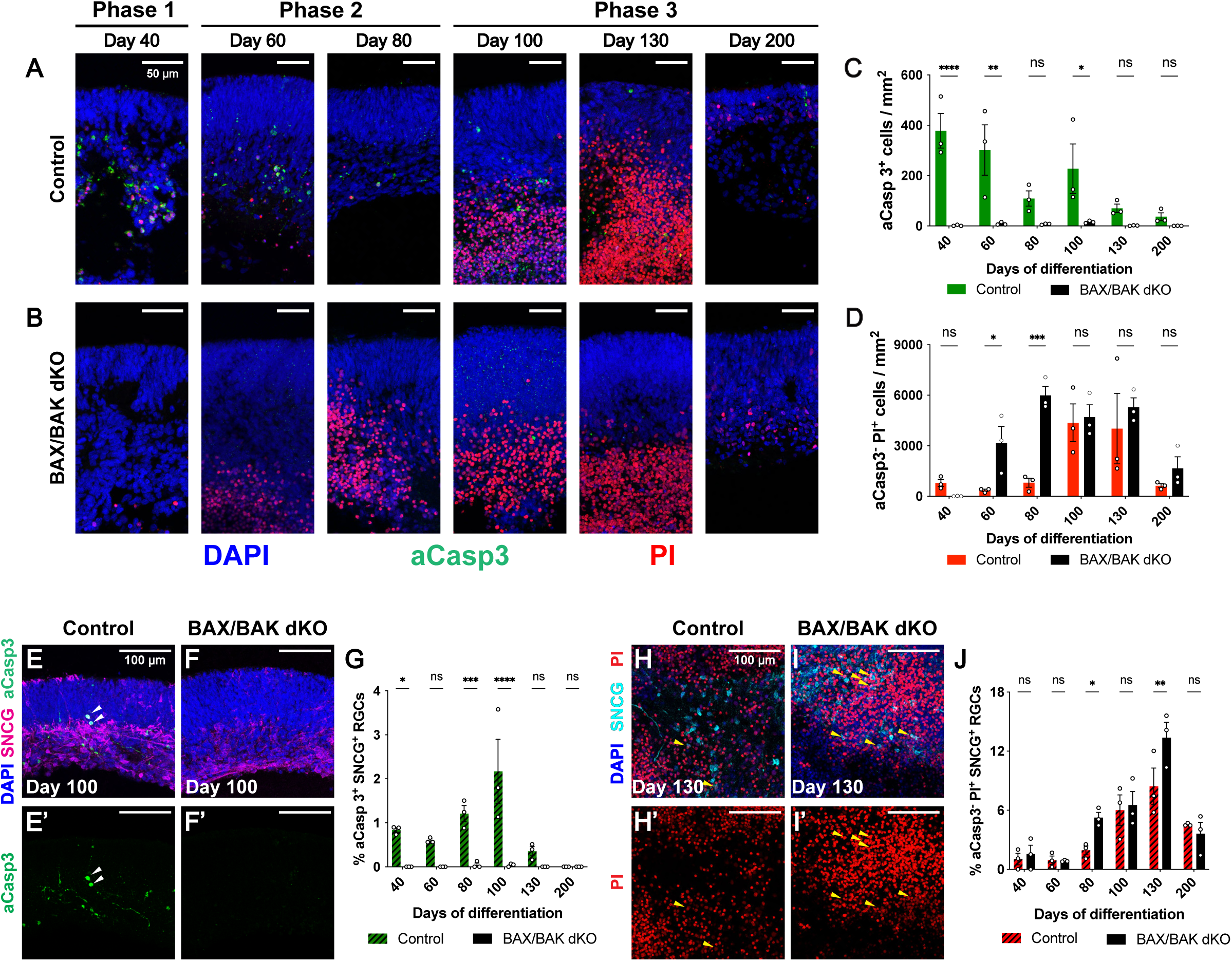
Apoptosis is blocked in *BAX/BAK* mutant organoids. **(A-B)** aCasp3 (green) and PI (red) in control (A) and *BAX/BAK dKO* (B) organoids. **(C-D)** Density of aCasp3^+^ apoptotic cells in retinal layers (C) and aCasp3^+^ PI^-^ necrotic cells in the core of control and *BAX/BAK dKO* organoids. Data are presented as mean ± SEM. N = 3 organoids per timepoint. Two-way ANOVA followed by Šidák’s post hoc test. * p < 0.05, ** p < 0.01, *** p < 0.001, **** p < 0.0001, ns = not significant. **(E-F)** aCasp3^+^ SNCG^+^ RGCs (white arrowheads) in control (E) and *BAX/BAK dKO* (F) retinal organoids. **(G)** % aCasp3^+^ SNCG^+^ RGCs of all SNCG^+^ RGCs in control and *BAX/BAK dKO* organoids. Data are presented as mean ± SEM. N = 3 organoids per timepoint. Data are presented as mean ± SEM. N = 3 organoids per timepoint. Two-way ANOVA followed by Šidák’s post hoc test. * p < 0.05, *** p < 0.001, **** p < 0.0001, ns = not significant. **(H-I)** PI^+^ SNCG^+^ RGCs (yellow arrowheads) in control (H) and *BAX/BAK dKO* (I) organoids. **(J)** % aCasp3^-^ PI^+^ SNCG^+^ RGCs of all SNCG^+^ RGCs in control and *BAX/BAK dKO* organoids. Data are presented as mean ± SEM. N = 3 organoids per timepoint. Two-way ANOVA followed by Šidák’s post hoc test. * p < 0.05, ** p < 0.01, ns = not significant.

To determine how *BAX/BAK dKO* affects necrosis in organoids, we measured the density of necrotic cells (Casp3^-^ PI^+^) in the core during differentiation (**Fig. 2A-B**). In control organoids, the density of necrotic cells was low in Phases 1 and 2 on days 40, 60, and 80, increased in Phase 3 on days 100 and 130, and then decreased on day 200 (**Fig. 2A, 2D**). In *BAX/BAK dKO* organoids, the density of necrotic cells was low at day 40 but increased during an extended wave of necrosis in Phases 2 and 3 from day 60 to day 130, followed by a decrease on day 200 (**Fig. 2B, 2D**). Necrotic cell density was significantly higher in *BAX/BAK dKO* organoids during Phase 2 on days 60 and 80 (**Fig. 2D**), suggesting that loss of apoptosis leads to earlier and more extensive necrosis.

SNCG (γ-synuclein) is highly expressed in developing and mature mammalian RGCs (Surguchov et al., 2001; Trimarchi et al., 2007; Surgucheva et al., 2008). To validate if *BAX/BAK dKO* abolishes apoptosis of RGCs, we measured the proportion of apoptotic RGCs (aCasp3^+^ SNCG^+^) among all SNCG^+^ RGCs (**Fig. 2E-G**). In control organoids, apoptosis of RGCs peaked on day 100 in early Phase 3 (**Fig. 2E, 2G**), similar to our observations in organoids derived from other cell lines (**Fig. 1L, S4F**). In contrast, apoptotic RGCs were nearly undetectable in *BAX/BAK dKO* organoids across all timepoints (**Fig. 2F-G**). Thus, apoptosis of RGCs is blocked in *BAX/BAK dKO* organoids.

To determine how *BAX/BAK dKO* affects necrosis of RGCs, we measured the proportion of necrotic RGCs (aCasp3^-^ PI^+^ SNCG^+^) in the core among all SNCG^+^ RGCs (**Fig. 2H-J**). In control organoids, necrotic RGCs were rare in Phases 1 and 2 (**Fig. 2J**). Their proportion increased in early Phase 3 on days 100 and 130 and decreased on day 200 (**Fig. 2H, 2J**). In *BAX/BAK dKO* organoids, the proportion of necrotic RGCs was low on days 40 and 60 and increased later, on days 80, 100 and 130, followed by a decline in late Phase 3 on day 200 (**Fig. 2I-J**). Necrosis of RGCs was elevated in *BAX/BAK* dKO organoids on days 80 and 130 (**Fig. 2J**), suggesting that blocking apoptosis leads to increased necrosis of RGCs.

Taken together, inhibition of BAX/BAK-mediated apoptosis blocks the two waves of apoptosis, yields increased retinal layer thickness, and increases necrosis during long-term organoid culture.

### Blocking apoptosis increases nRPC and RGC abundance in retinal organoids

To assess how blocking apoptosis affects retinal organoid development, we performed 10X Chromium snRNA-seq on control and *BAX/BAK dKO* organoids collected on days 50, 100, 150, and 200 (**Fig. 3A**). At each timepoint, four control and four *BAX/BAK dKO* organoids from the same batch were collected. After quality control and filtering, we profiled 29,821 nuclei in total. For dimensionality reduction, we applied non-negative matrix factorization (NMF) to identify gene expression patterns in our dataset (Stein-O’Brien et al., 2019; Johnson et al., 2023) (**Fig. S6A**). Because human retinal organoids can also generate retinal pigmented epithelium (RPE) cells and other non-retinal cell types (e.g., brain and spinal cord-like cells) (Liu et al., 2023), we manually annotated these populations using known markers and excluded them from downstream analysis (**Fig. S6B-D**).

**Figure 3.**
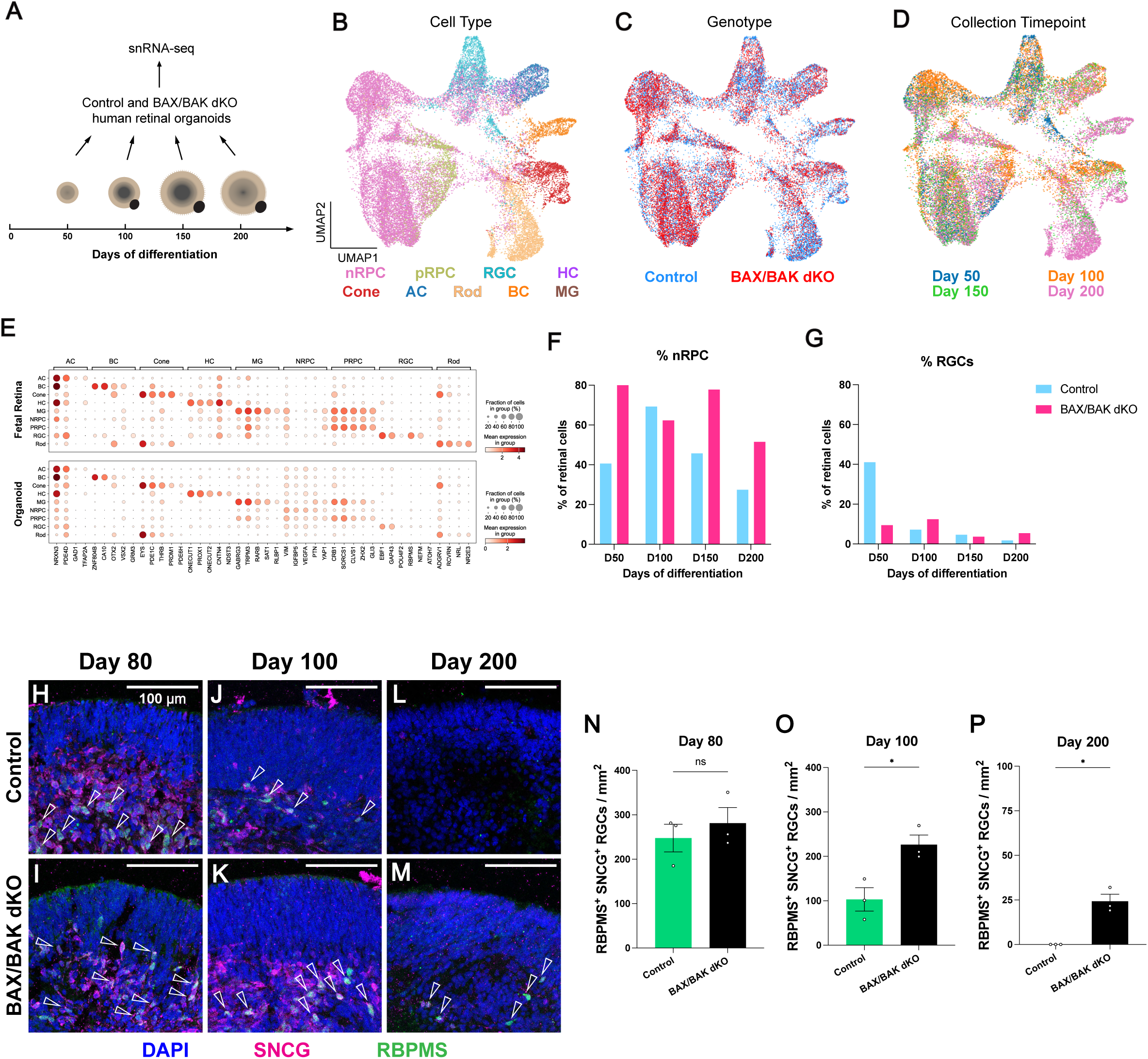
Blocking apoptosis leads to increases in nRPCs and RGCs in organoids. **(A)** Design of snRNA-seq experiments. Control and *BAX/BAK dKO* retinal organoids were collected on days 50, 100, 150 and 200. **(B-D)** Uniform manifold approximation and projection (UMAP) of retinal organoid-derived snRNA-seq data, colored by retinal cell types (B), genotypes (C), or collection timepoints (D). nRPC, neurogenic RPC; pRPC, proliferative RPC; RGC, retinal ganglion cell; HC, horizontal cell; Cone, cone photoreceptor; AC, amacrine cell; Rod, rod photoreceptor; BC, bipolar cells; MG, Muller glia. **(E)** Validation of marker genes for retinal cell types in the fetal retina reference dataset (upper) and retinal organoid dataset (lower). **(F-G)** % annotated nRPCs (E) and RGCs (F) of retinal cells at each collection timepoint. Blue = control; red = *BAX/BAK dKO*. **(H-P)** Maintenance of RBPMS^+^ SNCG^+^ RGCs (empty arrowheads) in control (H, J, L) and *BAX/BAK dKO* (I, K, M) organoids on days 80, 100, and 200. (N-P) RBPMS^+^ SNCG^+^ RGC density in retinal layers of control and *BAX/BAK dKO* organoids on days 80 (N), 100 (O), 200 (P). Data are presented as mean ± SEM. N = 3 organoids per timepoint. For N and O, unpaired two-tailed student’s t-test, * p < 0.05, ns = not significant; for P, unpaired two-tailed Welch’s t-test, * p < 0.05.

To annotate retinal cell types in our dataset, we trained a random forest classifier on a published single-nucleus dual-omics atlas of the human fetal retina as the reference (Breiman, 2001; Zuo et al., 2024) (**Fig. S6E-H**). Cell type annotations for organoid-derived cells were then reprojected for visualization (**Fig. 3B**). All major retinal cell types were present for control and *BAX/BAK dKO* organoids (**Fig. 3B-D, S6I, Supplementary Tables**). Organoid-derived and fetal retinal cells exhibited highly similar expression of canonical marker genes and classifier-derived markers that define retinal cell types (**Fig. 3E**) (Lu et al., 2020; Zuo et al., 2024), validating the cell type annotations of the organoid dataset.

To determine how blocking apoptosis affects the abundance of retinal cell types, we compared the proportions of cell types. As the early wave of apoptosis affects nRPCs and newborn neuronal precursors (**Fig. 1S, S4M-O**), we first compared the proportions of cells classified as nRPCs, which include dividing nRPCs and newborn precursors. *BAX/BAK* dKO organoids displayed higher proportions of nRPCs on days 50, 150 and 200 (**Fig. 3F**). In contrast, differences in the proportions of pRPCs were minimal (**Fig. S6I-L**). These results suggest that blocking apoptosis leads to an increase in nRPCs, consistent with our findings that the early wave of apoptosis affects nRPCs and newborn neuronal precursors (**Fig. S4M-O**).

As the late wave of apoptosis affects RGCs, we next compared the proportions of RGCs. Surprisingly, *BAX/BAK dKO* organoids displayed a lower percentage of RGCs on day 50 (**Fig. 3G**), suggesting that early RGC neurogenesis is delayed when apoptosis is blocked. Nevertheless, *BAX/BAK dKO* organoids exhibited a comparable or increased proportion of RGCs on days 100, 150 and 200 (**Fig. 3G**), suggesting that RGC abundance is increased in *BAX/BAK dKO* organoids. To validate these observations, we quantified RGC density in control and *BAX/BAK dKO* organoids on days 80, 100, and 200 of differentiation. RGCs were identified based on co-expression of RGC markers SNCG and RBPMS. SNCG^+^ RBMPS^+^ RGC density was similar in control and *BAX/BAK dKO* organoids in Phase 2 on day 80 (**Fig. 3H-I, 3N**). In contrast, the densities of SNCG^+^ RBMPS^+^ RGCs were significantly increased in *BAX/BAK dKO* organoids in Phase 3 on days 100 and 200 (**Fig. 3J-M, 3O-P**). Similarly, *BAX/BAK dKO* organoids displayed an increased density of POU4F1^+^ RGCs on day 200 (**Fig. S6M-O**). These data show that blocking apoptosis increases RGC abundance during long-term organoid culture.

Together, blocking apoptosis increases the abundance of nRPCs/neuronal precursors and RGCs, consistent with their death in the early and late waves of apoptosis, respectively.

### Blocking apoptosis delays early RGC neurogenesis and improves RGC survival in retinal organoids

Since apoptosis affects nRPCs/neuronal precursors and RGCs, the increased density of RGCs in Phase 3 *BAX/BAK dKO* organoids may result from increased RGC neurogenesis and/or improved RGC survival.

To test how blocking apoptosis affects RGC neurogenesis, we pulsed control and *BAX/BAK dKO* organoids with EdU for 24 hours on days 36, 56, 76, and 96, and measured the percentage of newborn EdU^+^ SNCG^+^ RGCs among all EdU^+^ cells 4 days later (**Fig. 4A**). The proportion of EdU^+^ SNCG^+^ RGCs was higher in control compared to *BAX/BAK dKO* organoids following pulses on day 36 (**Fig. 4B-D**) but not on day 56 (**Fig. 4D**), suggesting that blocking apoptosis delays early RGC neurogenesis. In contrast, both genotypes showed similarly low proportions when pulsed on days 76 and 96 (**Fig. 4D**), indicating that blocking apoptosis does not extend the time window of RGC neurogenesis in organoids. We validated this temporal pattern of RGC neurogenesis with a separate stem cell line (**Fig. S7A-C**). In addition, we found no evidence for cytotoxic effects of day 76 EdU treatment in Phase 2 organoids (**Fig. S7D-E**), indicating that the decreased EdU^+^ RGC proportion is not caused by EdU-induced cell death (Haskins et al., 2020).

**Figure 4.**
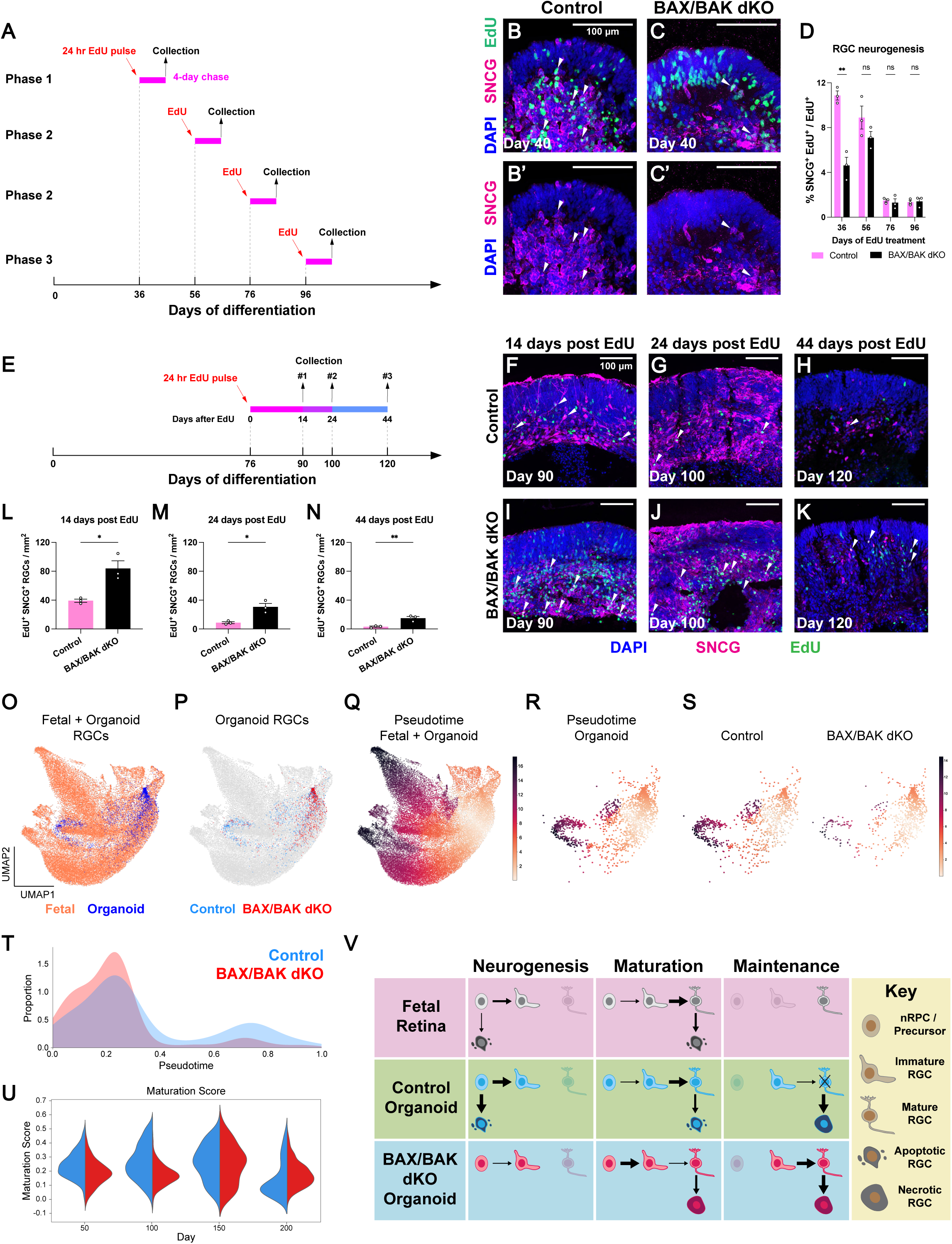
Blocking apoptosis delays RGC development and promotes RGC survival in organoids. **(A-D)** *BAX/BAK dKO* delays RGC neurogenesis. (A) Experimental design of RGC neurogenesis measurement by EdU labeling. (B-C) EdU^+^ SNCG^+^ RGCs (white arrowheads) in day 40 control (B) and *BAX/BAK dKO* (C) organoids. (D) % EdU^+^ SNCG^+^ RGC of all EdU^+^ cells in control and *BAX/BAK dKO* organoids following EdU treatment on days 36, 56, 76 and 96. Data are presented as mean ± SEM. N = 3 organoids per timepoint. Unpaired two-tailed student’s t-test at each timepoint, ** p < 0.01, ns = not significant. **(E-N)** *BAX/BAK dKO* promotes RGC survival. (E) Experimental design of RGC lifespan measurement by EdU labeling. (F-K) EdU^+^ SNCG^+^ RGCs (white arrowheads) in control (F-H) and *BAX/BAK dKO* (I-K) organoids 14, 24 and 44 days after EdU treatment. (L-N) Density of EdU^+^ SNCG^+^ RGCs 14 (L), 24 (M), and 44 (N) days after EdU treatment. Data are presented as mean ± SEM. N = 3 organoids per timepoint. Unpaired two-tailed student’s t-test at each timepoint, * p < 0.05, ** p < 0.01, ns = not significant. **(O-P)** Integrated UMAP maps of fetal and organoid-derived RGC snRNA-seq data colored by sample sources (O, orange = fetal retina, blue = organoid) or genotypes (P, blue = control organoid RGCs; red = *BAX/BAK dKO* organoid RGCs; gray = fetal retina RGCs). **(Q)** Integrated UMAP of fetal and organoid RGCs is colored by pseudotime. **(R-S)** UMAPs of all organoid RGCs (R), control organoid RGCs (S, left) and *BAX/BAK dKO* organoid RGCs (S, right) colored by pseudotime. **(T)** Proportional distribution of control (blue) and *BAX/BAK dKO* (red) organoid RGCs along the inferred pseudotime trajectory. **(U)** Violin plots of maturation scores of control (blue) and *BAX/BAK dKO* (red) organoid RGCs on days 50, 100, 150, and 200. **(V)** Model for cell death and RGC development in human fetal retinas and retinal organoids.

To test whether blocking apoptosis improves RGC survival, we pulsed control and *BAX/BAK dKO* organoids with EdU for 24 hours on day 76 and measured the densities of EdU^+^ SNCG^+^ RGCs 14, 24, or 44 days later (**Fig. 4E**). Control organoids displayed a near-complete loss of EdU^+^ SNCG^+^ RGCs by day 44 after the EdU pulse, suggesting that most RGCs born on day 76 have a lifespan shorter than 44 days (**Fig. 4F-H**). We validated the lifespan of RGCs with a separate stem cell line (**Fig. S7F-J**). In contrast, *BAX/BAK dKO* organoids displayed higher densities of EdU^+^ SNCG^+^ RGCs compared to the control organoids on days 14, 24, and 44 after the EdU pulse (**Fig. 4I-N**), suggesting that RGC survival is improved in *BAX/BAK dKO* organoids.

Taken together, blocking apoptosis delays early RGC neurogenesis and improves RGC survival in *BAX/BAK dKO* retinal organoids.

### Blocking apoptosis slows maturation but promotes survival of mature RGCs

We next investigated RGC maturation to understand how apoptosis regulates overall RGC development. To compare RGC development in human retinal organoids and fetal retinas, we used transfer learning to project annotated organoid-derived RGCs onto the UMAP of human fetal RGCs from the reference snRNA-seq dataset (**Fig. 4O-P**) (Stein-O’Brien et al., 2019; Zuo et al., 2024).

To evaluate the progression of RGC maturation, we performed pseudotime analysis on fetal and organoid-derived RGCs using scFates (**Fig. 4Q**) (Faure et al., 2023). The inferred pseudotime trajectory highly aligns with the progression of biological ages in human fetal RGCs (**Fig. S7K-M**), suggesting that the pseudotime trajectory approximated *in vivo* human RGC maturation. Human organoid-derived RGCs show similar pseudotime trajectories (**Fig. 4R**), validated by the expression of genes associated with different maturation stages of RGCs (*ATOH7, GFRA1, FGF12*) (**Fig. S7N**) (Brown et al., 2002; Kay et al., 2005; Brzezinski et al., 2012; Kretz et al., 2006; Wang et al., 2016).

To assess how blocking apoptosis affects the progression of RGC maturation, we compared the distribution of RGCs from *BAX/BAK dKO* and control organoids along the pseudotime trajectory (**Fig. 4S**). Control organoids showed a greater proportion of RGCs at later pseudotime, whereas *BAX/BAK dKO* organoids were enriched for RGCs at early pseudotime (**Fig. 4T**). These results suggest that blocking apoptosis impedes maturation of RGCs.

To determine how the transcriptome changes during human RGC maturation, we used scFates to identify a set of 142 genes associated with mid-late and late stages of RGC maturation from a human fetal retina dataset (**Fig. S7O, Supplementary Tables**). Expression of these genes generally correlated with pseudotime progression of RGCs in control and *BAX/BAK dKO* organoids (**Fig. S7P-Q**). To quantify the expression of these maturation-associated genes, we then calculated an aggregate gene expression score (“maturation score”) for the same gene set (Wolf et al., 2018). The maturation score increased with the biological ages of fetal-derived RGCs (**Fig. S7R**), suggesting that it captures the developmental progression of human fetal RGCs.

To assess how blocking apoptosis affects transcriptomic changes during RGC maturation, we compared maturation scores of RGCs from control and *BAX/BAK dKO* organoids on days 50, 100, 150 and 200. On days 50 and 100, *BAX/BAK dKO* RGCs showed lower maturation scores than control RGCs (**Fig. 4U**), consistent with delayed RGC neurogenesis and maturation upon inhibition of apoptosis (**Fig. 4B-D, 4T**). On day 150, maturation scores were comparable between the two genotypes (**Fig. 4U**). On day 200, *BAX/BAK dKO* RGCs exhibited higher maturation scores than control RGCs (**Fig. 4U**), suggesting that blocking apoptosis leads to improved survival of more mature RGCs, consistent with our immunofluorescent staining results (**Fig. 3H-P**).

Taken together, our data suggest that RGCs in control organoids are generated, mature, and die by 200 days of differentiation, while RGCs in *BAX/BAK dKO* organoids are generated and mature more slowly but survive more readily until day 200.

## Discussion

We characterize here the spatiotemporal dynamics of cell death in human fetal retinas and retinal organoids, develop a strategy to promote RGC long-term survival in organoids by blocking apoptosis, and reveal a regulatory role of developmental apoptosis in RGC development. Human fetal retinas exhibit two waves of developmental apoptosis, with the first wave affecting nRPCs and neuronal precursors, and the second wave targeting RGCs and other retinal neurons (**Fig. 4V**). Human retinal organoids similarly display two waves of apoptosis, as well as an additional wave of necrosis (**Fig. 4V**). When apoptosis is blocked in organoids, RGC neurogenesis and maturation are delayed, while their survival is improved (**Fig. 4V**). These findings provide new insights into the roles of apoptosis in human RGC development and unveil promising strategies to promote RGC long-term survival in organoid models for future therapeutic applications.

### Comparing developmental cell death between human retinas and retinal organoids

Our studies reveal key similarities and differences in developmental apoptosis between human retinas and organoids. The patterns of apoptosis are broadly similar. In both fetal retinas and retinal organoids, an early wave of apoptosis affects RPCs and neural precursors, followed by a late wave that selectively eliminates a subset of RGCs and other neurons. At the peaks of the late wave, ∼1.0% of RBPMS^+^ RGCs were apoptotic in human fetal retinas (**Fig. S1H**), whereas ∼4.4% of mNeon^+^ RGCs were apoptotic in organoids (**Fig. 1L**). Though apoptosis plays a central part in regulating RGC development in both contexts, necrosis emerges as an additional mechanism of cell death causing RGC loss during long-term organoid culture (**Fig. 1O**).

These results suggest that organoids may lack essential extrinsic factors required for sustained RGC survival and maturation, such as neurotrophic factors (Lom and Cohen-Cory, 1999), physiological electrical stimuli (Tian and Copenhagen, 2003; McLaughlin et al., 2003; Whitney et al., 2023), and proper oxygen supply (Osborne et al., 2004; Kaur et al., 2008). Further investigation will focus on recapitulating the supporting factors present in the human fetal environment during organoid culture.

### Apoptosis regulates developmental timing of RGC neurogenesis

Blocking the early wave of apoptosis of nRPCs and neural precursors leads to an increased population of nRPCs and delays in RGC neurogenesis and maturation, suggesting a non-cell-autonomous mechanism regulates RGC production. Our findings suggest that the early wave of apoptosis lowers the RPC-to-neuron ratio by eliminating nRPCs and neuronal precursors, promoting RGC neurogenesis and retinal development. Blocking apoptosis shifts the ratio towards nRPCs/neural precursors by increasing their relative abundance to neurons, which delays RGC neurogenesis and maturation. In the developing mouse retina, inhibition of apoptosis by *Bax* and *Bak* knockout does not increase cell proliferation (Hahn et al., 2003; Ke et al., 2018). Mouse RPC proliferation and differentiation are regulated by RGC numbers via the Shh signaling pathway, highlighting a critical role of RPC-to-neuron ratio in coordinating the timing of retinal development (Wang et al., 2005; Mu et al., 2005). Recently, we found a similar role for the RPC-to-neuron ratio in regulating the photoreceptor developmental timing via thyroid hormone levels and the dynamic expression in RPCs of DIO3, a thyroid hormone degrading enzyme (McNerney et al., 2025). It appears that the shifting RPC-to-neuron ratio plays an active role in regulating the timing of retinal neurogenesis and other developmental mechanisms.

### Promoting RGC survival in human retinal organoids

Different strategies have been applied to promote RGC survival in long-term organoid culture. Generation of assembloids by fusing human retinal organoids and brain organoids allows robust RGC neurite outgrowth, reduces apoptosis, and enhances the maintenance of RGCs through day 150. Similarly, providing brain-derived neurotrophic factor (BDNF) promotes RGC maintenance in organoids (Fligor et al., 2021). Generation of vascularized human retinal organoids by co-culturing growing retinal organoids and epithelial cells reduces apoptosis and promotes the upregulation of RGC markers (Inagaki et al., 2025). In our study, genetic knockout of *BAX* and *BAK* blocks RGC apoptosis and prolongs the survival of SNCG^+^ RBPMS^+^ RGCs through day 200 of differentiation (**Fig. 3H-P**), highlighting the contribution of apoptosis to the progressive loss of RGC in organoids.

Although human retinal organoids cultured in bioreactors or on microfluidic chips display increased size, reduced hypoxic stress, and increased yields of photoreceptors and RGCs (Ovando-Roche et al., 2018; Xue et al., 2022; Gong et al., 2023; Drabbe et al., 2025), how these strategies alter necrosis and long-term RGC survival remains unclear. Our study characterizes the formation of the necrotic core and how necrosis causes RGC loss (**Fig. 1M-O, S3C-E**), underscoring the need to reduce necrosis in long-term organoid culture to improve the yield of mature RGCs. Air-liquid interface culture of brain organoid slices alleviates interior hypoxia and facilitates their expansion and lamination, making it a promising assay for reducing necrosis in the core of organoids (Giandomenico et al., 2019; Qian et al., 2020; Guy et al., 2021). Future efforts will develop strategies combining inhibition of apoptosis with reduction of necrosis to further enhance RGC survival in retinal organoids, thereby advancing their translational potential for transplantation-based therapies of optic neuropathies (Soucy et al., 2023).

## Conclusion

Taken together, this study deepens our understanding of the mechanisms that regulate human RGC neurogenesis, maturation, and survival. Moving forward, advances in organoid engineering that minimize loss and promote maturation of RGCs will be crucial for improving disease modeling and developing regenerative cell therapies for neurodegenerative diseases of the human visual system.

**Figure S1.**
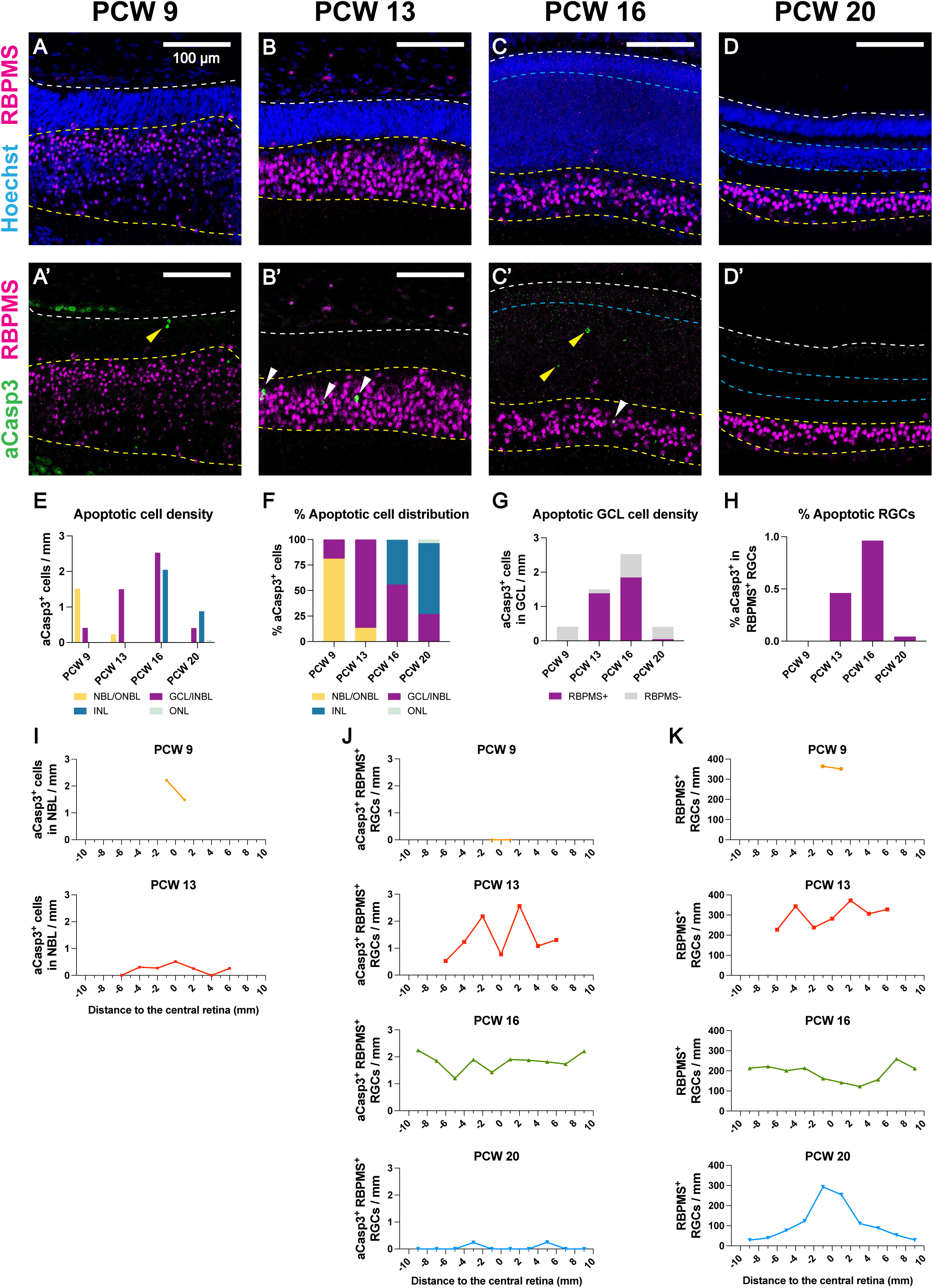
Two waves of apoptosis in developing human retinas. **(A-D)** Immunofluorescent staining of human fetal retina sections at post-conception week 9 (A), 13 (B), 16 (C) and 20 (D). The apical boundary of the neural retina (white dashed line) and the boundary of the INL (cyan dashed line) are outlined based on DAPI staining. The boundary of the GCL (yellow dashed line) is outlined based on RBPMS staining (magenta) in the GCL. White arrowheads indicate aCasp3^+^ RBPMS^+^ cells (apoptotic RGCs). Yellow arrowheads indicate aCasp3^+^ RBPMS^-^ cells (apoptotic non-RGCs). **(E-F)** Density (E) and laminar distribution (F) of apoptotic cells in layers of human fetal retinas at PCW 9, 13, 16 and 20. At PCW 9 and 13, the pseudostratified retinas are comprised of the NBL (or outer neuroblastic layer, ONBL) and the laminating GCL (or inner neuroblastic layer, INBL). At PCW 16 and 20, the fetal retinas are comprised of the ONL, INL and GCL. **(G)** Density of apoptotic RBPMS^+^ and RBPMS^-^ cells in the GCL of human fetal retinas at PCW 9, 13, 16 and 20. **(H)** Proportion of apoptotic RBPMS^+^ RGCs of all RBPMS^+^ RGCs in human fetal retinas at PCW 9, 13, 16 and 20. **(I)** Regional density of aCasp3^+^ cells in the NBL along the vertical meridian of human fetal retinas. **(J)** Regional density of aCasp3^+^ RBPMS^+^ RGCs along the vertical meridian of human fetal retinas. **(K)** Regional density of RBPMS^+^ RGCs along the vertical meridian of human fetal retinas.

**Figure S2.**
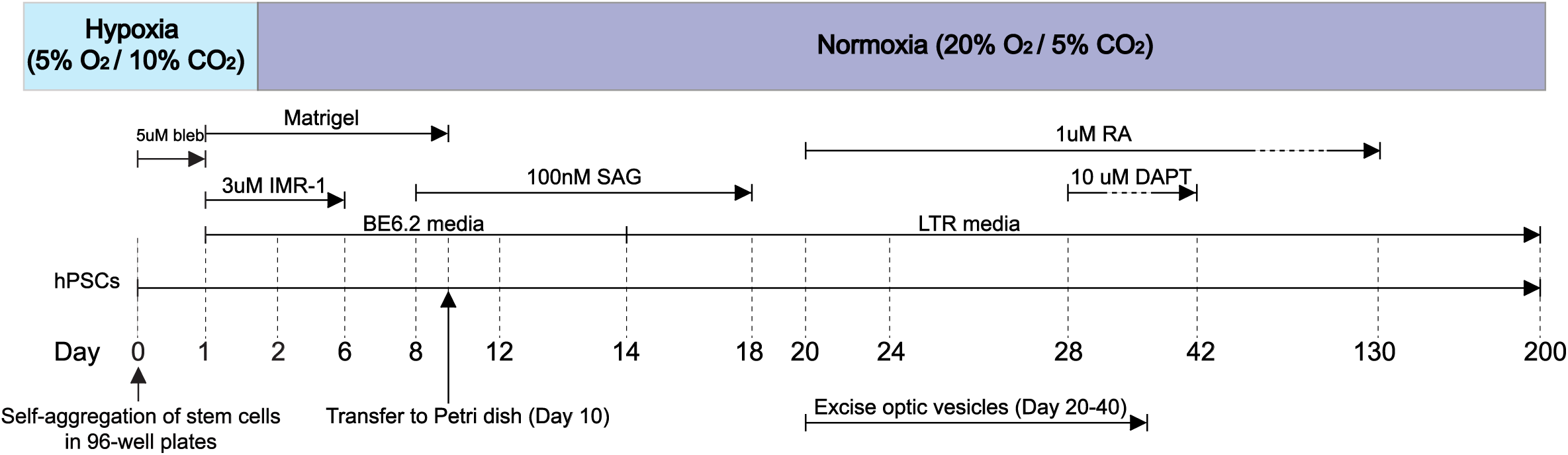
Gravity aggregation protocol of human retinal organoid differentiation. Gravity aggregation protocol adapted from previous studies (Wahlin et al., 2017; Eldred et al., 2018).

**Figure S3.**
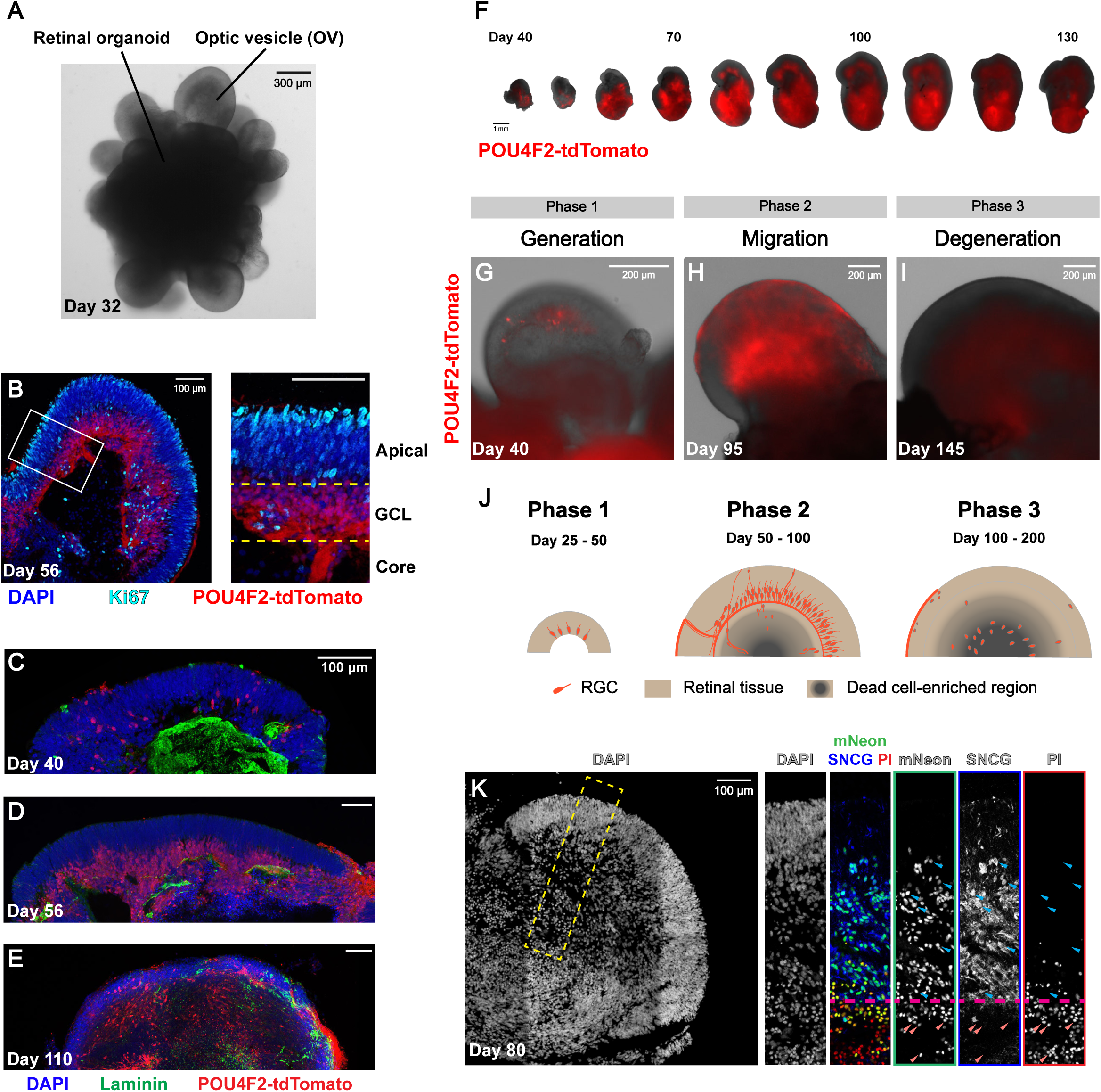
RGC migration and loss shown by RGC-specific reporters in human retinal organoids. **(A)** Differential interference contrast (DIC) image of a day 32 retinal organoid with OVs. **(B)** OV with lamination of the NBL, GCL, and core. (Inset) The boundaries of the NBL, GCL and the core (yellow dashed line) are indicated by Ki67^+^ RPCs (cyan) in the NBL and POU4F2-tdTomato^+^ RGCs (red) in the GCL. **(C-E)** ILM bleaching and RGC migration to the core on days 40 (C), 56 (D) and 110 (E). ILM is marked by laminin staining (green). **(F)** Overlaying DIC and red fluorescent images of a single H7 *POU4F2-tdTomato* retinal organoid from day 40 to day 130. **(G-I)** Overlaying DIC and fluorescent images of Phase 1 (G), Phase 2 (H), and Phase 3 (I) organoids. **(J)** Schematic of three phases of RGC development and loss in human retinal organoids. **(K)** mNeon labeled both live and dead RGCs in human retinal organoids. A region of interest is outlined and shown in the insets (yellow dashed line). (Insets) Live mNeon^+^ RGCs (SNCG^+^ PI^-^ mNeon^+^, blue arrowhead) express the RGC-specific marker SNCG and are predominantly located in the retinal layers. Dying and dead mNeon^+^ RGCs (SNCG^-^ PI^+^ mNeon^+^, orange arrowhead), labeled by propidium iodide (PI), are mostly located in the core. The boundary between the retinal layers and the core of OVs (magenta dashed line) is outlined based on the distribution of pyknotic nuclei and PI staining.

**Figure S4.**
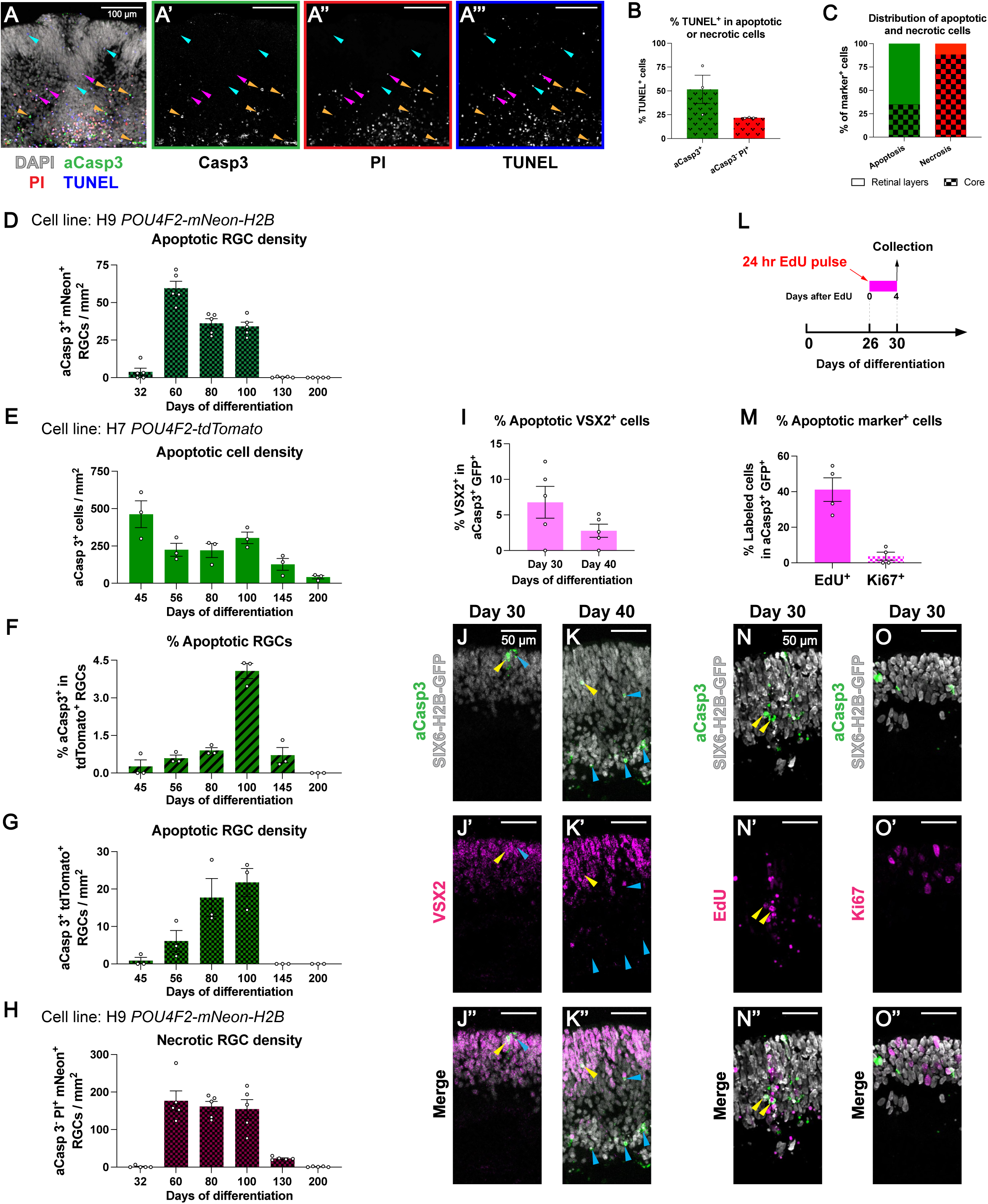
Measurements of apoptosis and necrosis in human retinal organoids. **(A)** Co-detection of dead cells with aCasp3, PI, and TUNEL staining in day 60 organoids derived from GM25256 hiPSCs. Orange arrowheads indicate TUNEL^+^ aCasp3^+^ cells (TUNEL^+^ apoptotic cells). Magenta arrowheads indicate TUNEL^+^ aCasp3^-^ PI^+^ cells (TUNEL^+^ necrotic cells). Cyan arrowheads indicate TUNEL^+^ aCasp3^-^ PI^-^ cells (TUNEL^+^ cells without apoptotic or necrotic markers). **(B)** % TUNEL^+^ cells of apoptotic or necrotic cells in day 60 retinal organoids. Data are presented as mean ± SEM. N = 3 organoids. **(C)** Distribution of apoptotic and necrotic cells in retinal layers (solid) and the core (checkerboard) of day 60 organoids. N = 3 organoids. **(D)** Density of aCasp3^+^ mNeon^+^ apoptotic RGCs in retinal layers of H9 *POU4F2-mNeon-H2B* retinal organoids. Data are presented as mean ± SEM. N = 5 organoids per timepoint. **(E-G)** Density of apoptotic cells (E), % apoptotic tdTomato^+^ RGCs (F), and density of apoptotic tdTomato^+^ RGCs (G) in H7 *POU4F2-tdTomato* retinal organoids. Data are presented as mean ± SEM. N = 3 organoids per timepoint. **(H)** Density of aCasp3^-^ PI^+^ mNeon^+^ necrotic RGCs in the core of H9 *POU4F2-mNeon-H2B* OVs. Data are presented as mean ± SEM. Data are presented as mean ± SEM. N = 5 organoids per timepoint. **(I-K)** VSX2^+^ RPCs in the aCasp3^+^ GFP^+^ population during the early wave of apoptosis. (I) % VSX2^+^ cells of all aCasp3^+^ GFP^+^ cells on days 30 and 40. Data are presented as mean ± SEM. N = 5 organoids per timepoint. (J-K) VSX2^+^ (yellow arrowheads) and VSX2^-^ (blue arrowheads) cells among the aCasp3^+^ GFP^+^ cells in day 30 (J) and day 40 (K) organoids. **(L-O)** Identification of cell types of the aCasp3^+^ GFP^+^ population during the early wave of apoptosis. (L) Experimental design of EdU labeling of early apoptotic cells. EdU was pulsed for 24 hours on day 26. EdU-treated organoids were collected 4 days later. (M) % aCasp3^+^ GFP^+^ labeled by EdU^+^ or Ki67^+^ of all aCasp3^+^ GFP^+^ cells. Data are presented as mean ± SEM. N = 4 organoids per timepoint. (N-O) EdU^+^ (yellow arrowhead, N) or Ki67^+^ cells (O) that are aCasp3^+^ GFP^+^ in day 30 organoids.

**Figure S5.**
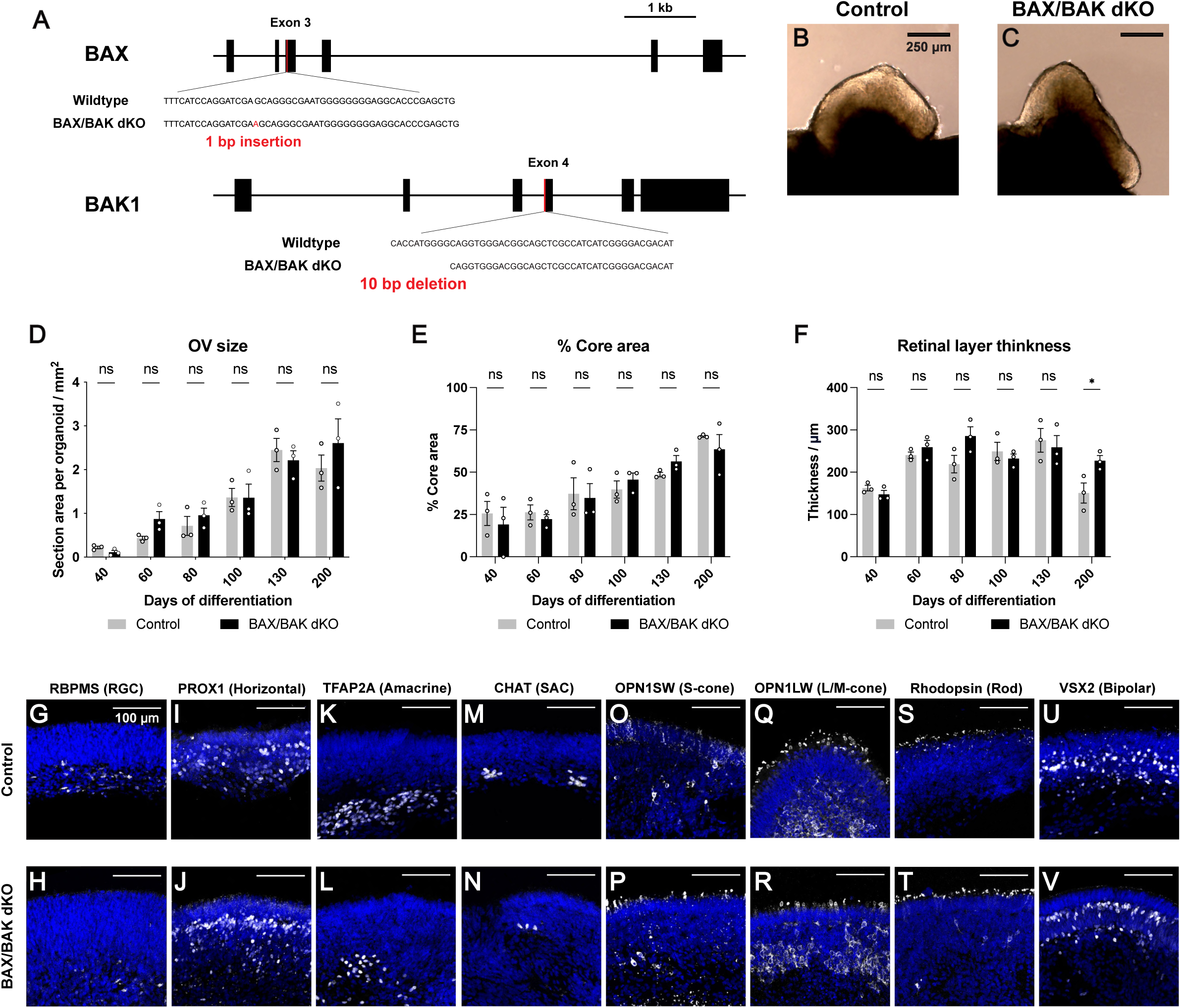
Development of human control and *BAX/BAK dKO* retinal organoids. **(A)** Schematic of mutations generated in the *BAX* and *BAK1* (*BAK*) loci (Joshi et al., 2020). Black bars indicate exons. Black lines between the bars indicate introns. Red bars indicate the location of mutations. **(B-C**) Brightfield images of OVs of day 25 control (B) and *BAX/BAK dKO* (C) retinal organoids. **(D-F)** Quantifications of OV size (D), % of core area (E), and thickness of retinal layers (F) in control and *BAX/BAK dKO* organoids. Data are presented as mean ± SEM. N = 3 organoids per timepoint. Two-way ANOVA followed by Šidák’s post hoc test, * p < 0.05, ns = not significant. **(G-V)** Major neuronal types are generated in control and *BAX/BAK dKO* retinal organoids, including RGCs (G-H, RBPMS^+^, day 100), horizontal cells (I-J, PROX1^+^, day 200), amacrine cells (K-L, TFAP2A^+^, day 200), starburst amacrines/SACs (M-N, CHAT^+^, day 200), S-cone photoreceptors (O-P, OPN1SW^+^, day 200), L/M-cone photoreceptors (Q-R, OPN1LW^+^, day 200), rod photoreceptors (S-T, Rhodopsin^+^, day 200), and bipolar cells (U-V, VSX2^+^, day 200).

**Figure S6.**
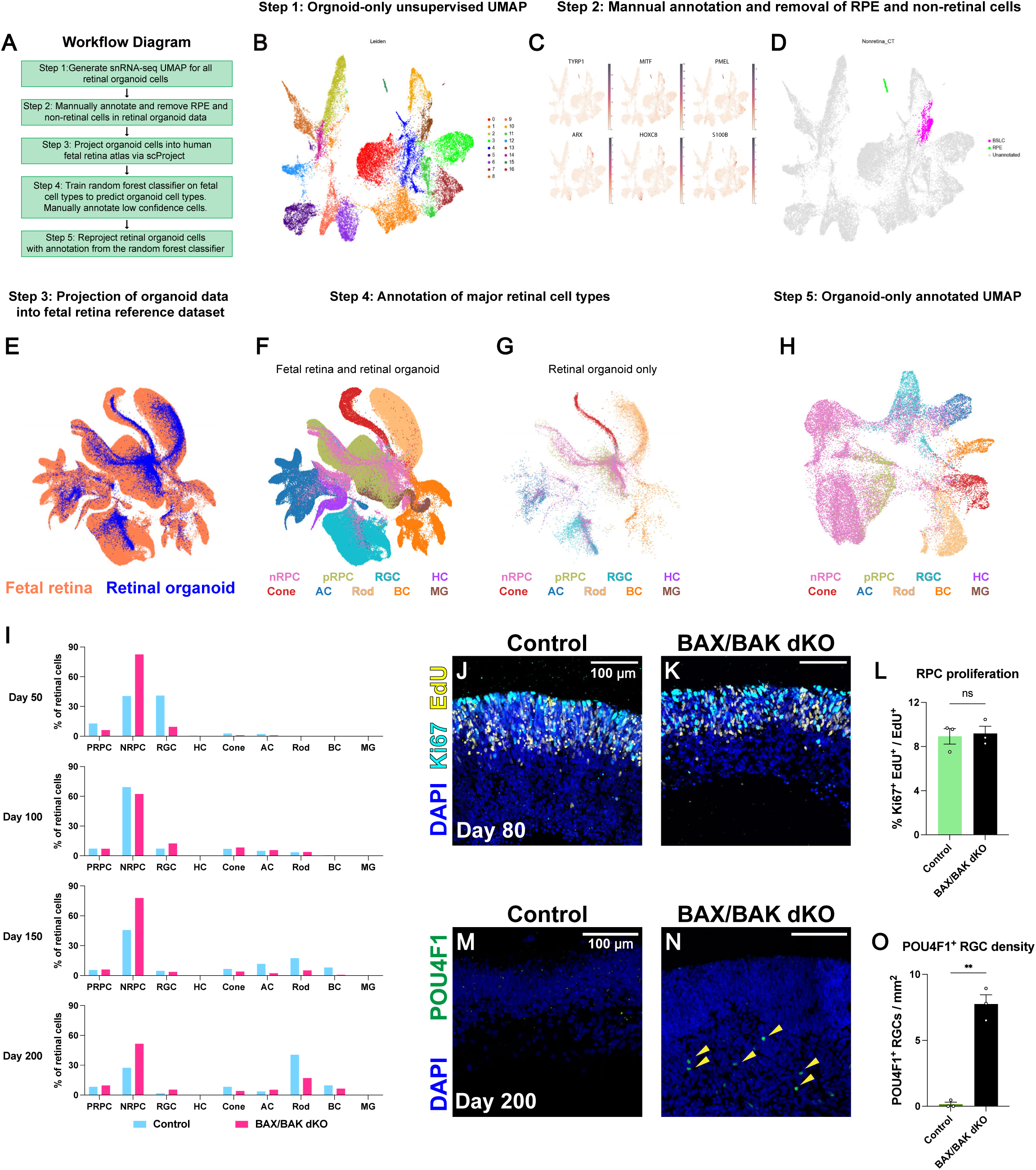
snRNA-seq analysis of control and *BAX/BAK dKO* organoids. **(A)** Diagram of major cell class annotation workflow. In steps 1-2, organoid-only data underwent dimension reduction and were projected onto UMAPs. RPE and non-retinal cells (brain and spinal cord-like cells, BSLCs) were manually annotated by known markers and removed from downstream analysis. In steps 3-4, organoid data were mapped to the reference fetal UMAP with scvi-tools and annotated by a random forest classifier trained on the fetal data. In step 5, annotated organoid-only data were reprojected. **(B)** Organoid-only data were projected and clustered onto a UMAP in an unsupervised manner. **(C-D)** RPE and non-retinal cells were manually annotated based on expression of known marker genes. (C) Expression of RPE and BSLC marker genes (*TYRP1*, *MITF*, *PMEL*, *ARX*, *HOXC8*, *S100B*). (D) RPE (green) and BSLC clusters (magenta) are highlighted in the unsupervised UMAP from step 1. (E) UMAP of integrated fetal retina and retinal organoid data from step 2. UMAP is colored by sample sources (fetal = orange, organoid = blue). **(F-G)** UMAPs of inferred major class annotation for development data from step 3. Integrated (F) and organoid-only UMAP (G) with annotation is shown. **(H)** UMAP of organoid-only data from step 4. At this step, fetal retina data were removed, and organoid-only data are reprojected. **(I)** % of retinal cell type abundance at each collection timepoint. Blue = control; red = *BAX/BAK dKO*. Difference in the proportions for cone photoreceptors, horizontal cells, amacrine cells, and Müller glia were minimal. Proportions of rod photoreceptors and bipolar cells were decreased in *BAX/BAK dKO* organoids on days 150 and 200, suggesting that blocking apoptosis impedes their neurogenesis. **(J-K)** Ki67^+^ EdU^+^ pRPCs in day 80 control (J) and *BAX/BAK dKO* (K) retinal organoids. EdU treatments for 24 hours were performed on day 76, and EdU-treated organoids were collected 4 days after each treatment. **(L)** % Ki67^+^ EdU^+^ cells of all EdU^+^ cells in day 80 control and *BAX/BAK dKO* organoids. Data are presented as mean ± SEM. N = 3 organoids. Unpaired two-tailed student’s t-test, ns = not significant. **(M-N)** POU4F1^+^ RGCs (green, yellow arrowhead) in day 200 control (M) and *BAX/BAK dKO* (N) organoids. **(O)** Density of POU4F1^+^ RGCs in day 200 control and *BAX/BAK dKO* organoids. Data are presented as mean ± SEM. N = 3 organoids. Unpaired two-tailed Welch’s t-test, ** p < 0.01.

**Figure S7.**
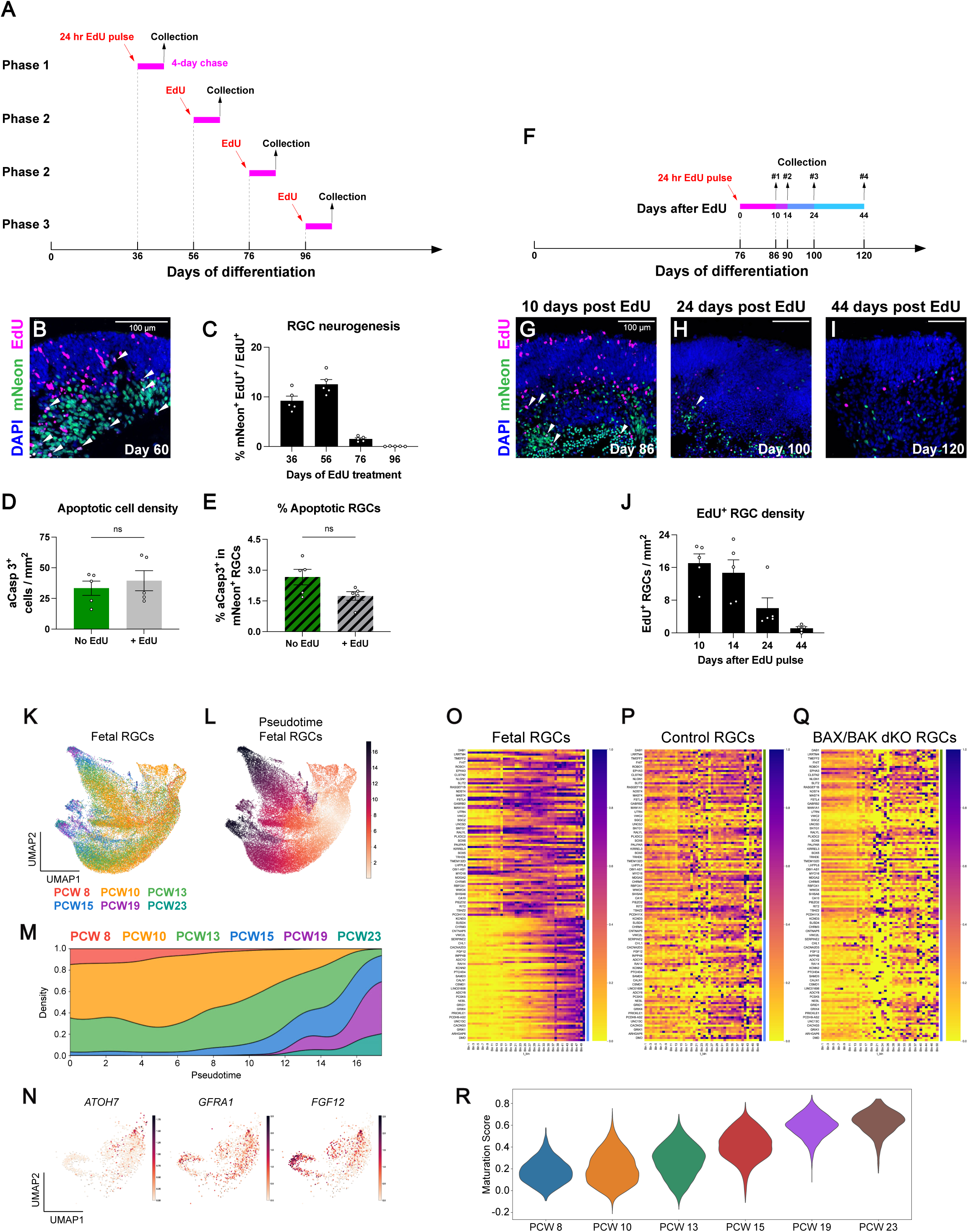
RGC neurogenesis, survival, and maturation in human control and *BAX/BAK dKO* retinal organoids. **(A)** Experimental design of RGC neurogenesis measurement by EdU labeling. EdU was pulsed for 24 hours on days 36, 56, 76 or 96. EdU-treated organoids were collected 4 days after each treatment. **(B)** EdU^+^ mNeon^+^ RGCs (white arrowhead) mark the RGC population born during the EdU treatment. **(C)** % newborn EdU^+^ mNeon^+^ RGCs of all EdU^+^ cells. Data are presented as mean ± SEM. N = 5 organoids per timepoint. (**D-E**) Density of apoptotic cells (D) and % apoptotic RGCs of all mNeon^+^ RGCs (E) in day 80 untreated (no EdU) and EdU-treated (EdU treatment at day 76) H9 *POU4F2-mNeon-H2B* retinal organoids. Data are presented as mean ± SEM. N = 5 organoids per timepoint. Unpaired two-tailed student’s t-test, ns = not significant. (F) Experimental design of RGC lifespan measurement by EdU labeling. EdU was pulsed for 24 hours on day 76. EdU-treated H9 *POU4F2-mNeon-H2b* organoids were collected 10, 14, 24 and 44 days after the treatment. **(G-I)** EdU^+^ mNeon^+^ RGCs (white arrowheads) 10 (G), 24 (H) and 44 (I) days after the EdU treatment. **(J)** Density of EdU^+^ mNeon^+^ RGCs after EdU treatment. Data are presented as mean ± SEM. N = 5 organoids per timepoint. **(K-L)** Fetal-derived RGC UMAPs colored by fetal age (K) and pseudotime (L). **(M)** Stacked density distribution of human fetal RGC data from the reference dataset along the pseudotime trajectory. Colors reflect different fetal ages of RGCs in the reference dataset. **(N)** Organoid-derived RGC UMAPs of representative genes in different RGC developmental stages (*ATOH7* = early*, GFRA1* = middle*, FGF12* = late) colored by expression level. **(O-Q)** Heatmaps of maturation-associated gene expression in the reference fetal (O), control organoid (P) and *BAX/BAK dKO* organoid (Q) RGCs binned by pseudotime units. Color blocks indicate genes associated with mid-late (green) and late (blue) stages of RGC maturation. **(R)** Violin plot of RGC maturation scores of human fetal RGCs in different PCWs.

## Materials and Methods

### Cell line maintenance

H7 *POU4F2-P2A-tdTomato-P2A-mTHY1.2* (modified H7 (WA07) human embryonic stem cells (hESCs), gift from the Zack lab at JHMI), H9 *POU4F2-P2A-mNeon-GFP* (modified H9 (WA09) hESCs, gift from the Wahlin lab at UCSD), IMR90.4 *SIX6-P2A-H2B-GFP* (modified hiPSCs, gift from the Zack lab at JHMI), wildtype GM25256 and *BAX/BAK dKO* GM25256 cells (modified hiPSCs, gift from the Gama lab at Vanderbilt University) were used for differentiation. Stem cells were maintained in mTeSR™1 (Cat# 85857, StemCell Technologies) on 1% (v/v) Matrigel-GFR™ (Cat# 354230, BD Biosciences) coated dishes and grown at 37°C and with 10% CO_2_ + 5% O_2_ in a HERAcell 150i or 160i incubator (Thermo Fisher Scientific). Cells were passaged every 4-5 days according to confluence as in previous studies (Wahlin et al., 2017). Cells were passaged with Accutase (Cat# SCR005, Sigma) for 8-12 minutes and dissociated to single cells. Cells in Accutase were added 1:2 to mTeSR™1 with 5 μM Blebbistatin (Bleb, Cat# B0560, Sigma), and pelleted at 150X gravity for 5 minutes. Cells were resuspended in mTeSR™1 with 5 μM Blebbistatin, density was quantified, and they were plated at 5,000-15,000 cells per well in a prepared 6-well Matrigel-coated plate. Cells were fed with mTeSR™1 48 hours after passaging and every subsequent 24 hours until passaged again. To minimize cell stress, no antibiotics were used. Cell lines were tested monthly for mycoplasma using MycoAlert (LT07, Lonza).

### Cell culture media

#### E6 supplement

970 μg/mL Insulin (Cat# 11376497001, Roche), 535 μg/mL holo-transferrin (Cat# T0665, Sigma), 3.20 mg/mL L-ascorbic acid (Cat# A8960, Sigma), 0.7 μg/mL sodium selenite (Cat# S5261, Sigma).

#### BE6.2 media

2.5% E6 supplement (above), 2% B27 supplement (50X) minus Vitamin A (Cat# 12587010, Gibco), 1% Glutamax (Cat# 35050061, Gibco), 1% NEAA (Cat# 11140050, Gibco), 1 mM Sodium Pyruvate (Cat# 11360070, Gibco), and 0.87 mg/mL NaCl in DMEM (Cat# 11885084, Gibco).

#### LTR (Long-term retina) media

25% F12 (Cat# 11765062, Gibco), 2% B27 supplement (50X) (Cat# 17504044, Gibco), 10% heat-inactivated FBS (Cat# 16140071, Gibco), 1mM Sodium Pyruvate (Cat# 11360070, Gibco), 1% NEAA (Cat# 11140050, Gibco), 1% Glutamax (Cat# 35050061, Gibco), and 1 mM taurine (Cat# T-8691, Sigma) in DMEM (Cat# 11885084, Gibco).

### Differentiation of human retinal organoids

Retinal organoids were differentiated from H7 ESCs as described in Wahlin et al., 2017 with minor modifications (**Fig. S2**). Briefly, human pluripotent stem cells were well-maintained, and only cultures with minimal to no spontaneous differentiation were used for aggregation. To aggregate, cells were passaged in Accutase at 37°C for 12 minutes to ensure complete single-cell dissociation. Cells were seeded in 50 uL of mTeSR™1 at 3,000 cells/well into 96-well ultra-low adhesion round-bottom Lipidure coated plates (Cat# 51011610, NOF). Cells were placed in hypoxic conditions (10% CO_2_ and 5% O_2_) for 24 hours to enhance survival. Cells naturally aggregated by gravity over 24 hours.

On day 1, cells were moved to normoxic conditions (5% CO_2_). On days 1-3, 50 μL of BE6.2 media containing 3 μM Wnt inhibitor (IWR1e; 681669, EMD Millipore) and 1% (v/v) Matrigel were added to each well. Between days 4-9, 100 μL of media were removed from each well, and 100 uL of media were replenished. On days 4-5, 100 μL of BE6.2 media containing 3 μM Wnt inhibitor and 1% Matrigel was added. On days 6-7, 100 μL of BE6.2 media containing 1% Matrigel was added. On days 8-9, 100 uL of BE6.2 media containing 1% Matrigel and 100 nM Smoothened agonist (SAG; Cat# 566660, EMD Millipore) was added.

On day 10, cell aggregates (organoids) were transferred to 15 mL tubes, rinsed 2-3X in DMEM (Cat# 11885084, Gibco), and resuspended in BE6.2 media with 100 nM SAG in untreated 10 cm polystyrene petri dishes. From this point on, media was changed every other day. Organoids were monitored daily and manually separated if stuck together or to the plate.

On days 13-18, LTR media with 100 nM SAG was added.

On days 16-20, organoids were maintained in LTR, and washed 2X with 5 mL of DMEM before being transferred to new plates to wash off dead cells. To increase the yield of retinal tissue, 1 μM all-trans retinoic acid (ATRA; R2625, Sigma) was added to LTR medium from days 20-130 in all conditions. 10 μM Gamma-secretase inhibitor (DAPT; Cat# 565770, EMD Millipore) was added to LTR from days 28-42. Organoids were grown at low density (10-20 per 10 cm dish) to reduce aggregation.

Between days 20 and 40, optic vesicles-like structures (OVs) were manually dissected if needed using sharpened tungsten needles and collected. After dissection, organoids were transferred into 15 mL tubes and washed 2X with 3 mL of DMEM. After day 40, organoids were regularly culled if they failed to differentiate or maintain retinal tissue properly.

### Phase quantification in retinal organoids

Human retinal organoids between days 30 and 200 of differentiation were kept in petri dishes with organoid culture media and live imaged with the EVOS XL Core Cell imaging system before collection. Differential interference contrast (DIC) and fluorescent images (555 nm channel) of developing organoids were overlayed by the default function of the EVOS XL Core imaging system and used for manual determination of RGC developmental phases based on RGC abundance and distribution. For quantification of RGC developmental phase progression, individual organoids were transferred and kept growing in designated wells of a 24-well plate separately between days 40 and 130 of differentiation. DIC and fluorescent images of these organoids were taken every 10 days, and their RGC developmental phases were evaluated as mentioned above.

### Mycoplasma monitoring

Cell lines and organoid media were tested monthly for mycoplasma using MycoAlert (LT07, Lonza) and excluded if positive.

### Sample acquisition and cryosection

Human fetal retina samples were obtained from the laboratory of Ya-wen Chen at the Icahn School of Medicine at Mount Sinai and shipped in cold fixative. The fetal age of the donor was calculated based on clinical judgement. All fetal ages were converted to post-conception weeks in this study.

For cryosectioning, fetal retinas and organoids were fixed by 4% formaldehyde for 1 hour at room temperature (20-25°C), treated with a serial wash of 5% (3X quick rinse), 6.25% (for 30 minutes), 12.5% sucrose (for 30 minutes) in 0.1M PBS, and then left in 25% sucrose in 0.1M PBS at 4°C overnight. On the next day, human fetal retinas and organoids were flash frozen in OCT on dry ice and moved to -80°C for long-term storage. On the day of cryosectioning, samples were retrieved from -80°C to -20°C freezers to adapt to the new temperature for at least 30 minutes. Cryosectioning was performed on CryoStar NX50 (epredia). Fetal retinas were oriented and cryosectioned along the nasal-temporal axis at 12-16 μm. Organoids were cryosectioned at 10-12 μm. Sections were left to air-dry at room temperature for at least 3 hours, and then stored long-term at -80°C.

### Immunohistochemistry

Sections of human fetal retinas and organoids were retrieved from -80°C freezers, adapted to room temperature for 10 minutes, and then heated in HybEZ II Oven (ACDBio) at 50°C for 10-15 minutes to minimize remaining water droplets. Sections were washed quickly in 1X PBS to remove remaining OCT, then blocked and permeabilized in 10% donkey serum + 0.5% Triton-X 100 in PBS (blocking solution) for 1 hour at room temperature. Sections were then incubated with primary antibodies (**Supplementary Tables**) in blocking solution for 16-20 hours or overnight at 4°C. Sections were washed 3X for 15 minutes in PBS, and incubated with secondary antibodies (Thermo Fisher Scientific, donkey secondary antibodies conjugated to Alexa Fluor 488, 555 or 647) diluted 1:800 in blocking solution for 1 hour (2 hours for fetal retina sections) at room temperature. Sections were then washed 3X in PBS quickly to get ready for nuclei counterstaining. For organoid samples, sections were incubated in 6 μM DAPI (4’,6-diamidino-2-phenylindole) in 1X PBS for 4 minutes at room temperature. For fetal retina samples, sections were incubated in 1:2000 dilution of Hoechst (Cat# 40046, Biotium) in PBS for 10 minutes at room temperature. Sections were then washed 3X for 15 min in PBS and mounted for imaging in SlowFade Gold Antifade mountant (Cat# S36940, Thermo Fisher Scientific).

### Apoptosis and necrosis co-detection

Human retinal organoids were collected and put in 3.5 cm petri dishes (Cat# 229638, CELLTREAT) with organoid culture media. Propidium iodide (Cat# P4170, Sigma-Aldrich) was filter sterilized and diluted in culture media to the final concentration of 25 μg/mL. Organoids were incubated in propidium iodide-added organoid culture media for 12 hours or overnight and collected on the next day. Treated organoids were then fixed, frozen, and cryosectioned for immunohistochemistry as mentioned above.

To distinguish apoptotic and necrotic cells in organoid sections, an antibody against cleaved/active Caspase 3 (Cat# 9961, Cell Signaling) was used for immunofluorescent staining. For quantitative analysis, aCasp3^+^ PI^-^ cells were counted as early apoptotic cells, and aCasp3^+^ PI^+^ cells were counted as late apoptotic cells, while aCasp3^-^ PI^+^ cells were counted as necrotic cells. Only apoptotic cells located in the retinal layers and necrotic cells located in the core were quantified in longitudinal analyses.

### TUNEL assay

The Click-iT™ Plus TUNEL Assay (Cat# C10617, Invitrogen) was conducted on human retinal organoid sections based on the manufacturer’s instructions for tissue sections. Before the TUNEL assay, the sections were washed with 1X PBS for 3 times to remove remaining OCT. After the TUNEL assay, sections were handled following the immunohistochemistry protocol for primary and secondary antibody staining, starting from blocking and permeabilization as mentioned above, but protected from light to minimize photobleaching.

### EdU treatment and detection

EdU solution (Click-iT EdU Assay, Invitrogen) was diluted in DMSO and added to the organoid culture media to a working concentration of 10 μM for 24 hours in the incubator, before treated organoids were washed and transferred to EdU-free organoid culture media. Optimal concentration of EdU was adjusted based on the viability of cell lines and kept consistent within the same set of experiments.

The Click-iT Plus EdU Assay (Cat# C10640, Invitrogen) was conducted on human retinal organoid sections based on the manufacturer’s instructions for tissue sections. Before the EdU assay, sections were washed with 1X PBS for 3 times to remove remaining OCT. After the assay, sections were stained by primary and secondary antibodies following the immunohistochemistry protocol, starting from blocking and permeabilization as mentioned above, but with caution to minimize exposure to light and avoid photobleaching.

### Microscopy and image processing

Brightfield images were acquired with an EVOS XL Core Cell imaging system. Differential Interference Contrast images were acquired with an EVOS M500 Imaging system on DIC with a 4x or a 10x air objective lens (Thermo Fisher). Fluorescent images were acquired using a Zeiss LSM 980 laser scanning confocal fluorescent microscopy. Confocal images were acquired with similar settings for laser power, photomultiplier gain and offset, and pinhole diameter. Camera images were acquired at similar epifluorescence intensity and exposure. Sections were imaged at 8-22 optical sections and 1-2 μm step size. Fluorescent images with a maximum intensity projection were used for downstream analysis.

### Measurements and quantification

Manual counting and semi-automated quantification were conducted to calculate cell density using the Cell Counter and Analyze Particles function panels in ImageJ. GraphPad Prism was used for statistical analysis. All error bars represent SEM. Organoids with non-retinal regions larger than 70% of the section area were removed from analysis and assumed to not have properly differentiated.

For human fetal data, N = 1 human fetal eye sample per timepoint was quantified. For each fetal eye, 2 interleaved sections within 100 μm to each other from a series of parasagittal sections around the central parts of the eye were quantified and averaged for a better representation of quantitative measurement. For regionalized density quantification, images of each retinal section were split into non-overlapping 2 mm segments based on their spatial distribution along the dorsal-ventral axis. Cell densities of each segment were individually quantified. Measurements of cell densities were plotted with the central segment centered. For peak detection, a parametric hypothesis test for unimodality using quadratic regression was conducted for regional cell densities at each timepoint and a peak was called when p < 0.05.

For organoid data, N = 3-5 organoids from the same genetic background or experimental treatments per timepoint were quantified. For each organoid, 3 interleaved sections with 50 μm intervals to each other were quantified and averaged for a better representation of quantitative measurement. For compartmentalized density quantification, boundaries between retinal layers and the core on organoid sections were manually outlined based on the enrichment of pyknotic cells by DAPI and/or PI staining. The areas of retinal layers or the core were quantified using the Measure functions in ImageJ based on the regional outlines.

For density measurement, apoptotic cell densities were calculated by dividing the number of apoptotic cells in retinal layers by the area of retinal layers. Necrotic cell densities were calculated by dividing the numbers of necrotic cells in the core by the area of the core. SNCG^+^ RBPMS^+^ cell densities were calculated by dividing the numbers of SNCG^+^ RBPMS^+^ cells in retinal layers by the area of retinal layers. Other density measurements were calculated by dividing the numbers of labeled cells by the area of entire OV sections.

### Sequencing sample collection and nuclei lysis

4 control and 4 *BAX/BAK dKO* organoids with OVs were collected at each collection timepoint (days 50, 100, 150, and 200). Media was aspirated from the organoids and the tissue was submerged in liquid nitrogen to flash freeze. Organoid samples were stored at -80°C until library preparation, to reduce batch effect across samples. Nuclei were isolated from retinal organoids using the chilled lysis buffer (10 mM Tris-HCl, 10 mM NaCl, 3 mM MgCl_2_, 0.01% Tween-20, 0.01% Nonidet P-40, 0.001% Digitonin, 1mM DTT, 1% BSA). The homogenate was incubated on ice for 10 minutes, washed twice with a resuspension buffer (10 mM Tris-HCl, 10 mM NaCl, 3 mM MgCl_2_, 0.1% Tween-20, 1mM DTT, 1% BSA) and filtered through a series of 70 μm and 40 μm Flowmi filters. The nuclei were resuspended in an appropriate volume of diluted nuclei buffer (10X Genomics) to achieve a concentration of approximately 2000–5000 nuclei/μL. Nuclei concentration was determined using trypan blue and DAPI staining.

### Library preparation and preprocessing

Single-nucleus multiomics was performed following the CG000338 library construction protocol from 10x Genomics. Libraries were sequenced on an Illumina Novaseq 6000 through Psomagen sequencing services. Reads were aligned to the GRCh38 human reference genome with CellRanger 8.0, and the resulting cell-by-gene count matrix was processed with the scanpy package (Wolf et al., 2018).

Cells were filtered to exclude those with more than 14% of counts mapped to mitochondrial genes, less than 200 genes, or more than 30000 unique molecular identifiers (UMIs). Cells from individual datasets were merged, normalized to total read counts, and log transformed with the percentage of mitochondrial counts and ribosomal counts used as variables for the regress_out function. Count matrixes were scaled by depth and log normalized. Nonnegative Matrix Factorization (NMF, sklearn, n_components = 20) was used for dimensionality reduction. n_components for NMF was selected based on a PCA of the data. Leiden clusters were calculated and UMAP coordinates were generated with the leiden and umap tools. Leiden clusters representing non-retinal cell types, including retinal pigmented epithelium (RPE) and brain and spinal cord-like cells (BSLCs), were manually annotated by their high expression of marker genes prior to downstream analyses (Lu et al., 2020; Liu et al., 2023).

### Cell type annotation and transfer learning

To call retinal cell types, we trained a random forest classifier on a reference snRNA-seq dataset of human fetal retinas (Zuo et al., 2024). The top 50 differentially expressed genes between major retinal cell types in the fetal dataset were selected using scanpy’s rank_genes_groups tool and used to train the classifier (sklearn RandomForestClassifier, n_estimators = 200, max_features = None). The organoid dataset was subset to these genes for classification and organoid cells were assigned their highest probability cell type by the trained classifier. For organoid cells with low log-likelihood, probability scores or BSLC/RPE-like expression patterns, previous manual annotations were used. Annotated non-retinal cells were removed from downstream analyses.

To validate the retinal cell type annotation by the classifier, NMF (n_components = 40) was conducted on the top 3000 highly variable genes in the reference fetal dataset. To visualize organoid cells in the same space as fetal cells in the reference dataset, fetal and organoid cells were projected onto these NMF components using scProject’s Elastic_Net tool (alpha = 0.005, L1 = 0.005) (Stein-O’Brien et al., 2019).

To visualize annotated organoid cells only, we subset the count matrix of the organoid dataset to a collection of the top 50 differentially expressed genes from each major cell type in the fetal reference dataset and 43 cell cycle-associated genes. NMF (n_components = 20) was conducted on the subset matrix for dimensionality reduction, and UMAP coordinates of reprojected organoid cells were generated with the umap tool.

Percent abundances of annotated organoid cell types at each timepoint were calculated by a custom script (SZ_percabun_annotated.py), and significance was assessed by chi-squared tests for proportions with Benjamini-Hochberg, multiple testing corrections.

### Pseudotime analysis of RGCs

To quantify developmental progression along a cell fate trajectory, RGCs from the fetal and organoid datasets were subset. Organoid RGCs with low confidence scores from the classifier were excluded from the analysis to ensure rigor. Organoid RGCs were projected onto a fetal RGC UMAP using the same transfer learning pipeline described above. Fetal and organoid RGCs were assigned a pseudotime value using the scFates package’s tree tool, with fetal RGC precursors being selected as the root (Faure et al., 2023). To assess for significant changes in pseudotime progression in organoid RGCs, we used a permutation test. Genes associated with early, mid and late RGC fate were identified using scFates test_association tool. 5 clusters of pseudotemporally associated genes were identified, with cluster 2 and 3 corresponding to the mid-late and late maturation stages of RGCs, respectively. To evaluate maturation of RGCs by aggregate gene expression, genes from clusters 2 and 3 were used as targets for scanpy’s score_genes function to generate an RGC maturation score (Wolf et al., 2018) (**Supplementary Tables).** To assess for significant changes in maturity score between timepoints, permutation tests were used with Benjamini-Hochberg multiple testing correction.

### Data availability

All raw sequencing data generated in this study can be accessed at GEO accession number GSE305194. This paper analyzes existing, publicly available data (Zuo et al., 2024), accessible at CZ CELLxGENE Discover with accession code 5900dda8-2dc3-4770-b604084eac1c2c82. All analysis code used in this study can be accessed with GitHub repository https://github.com/BGuy2/SZ_BAX-BAKdKO/. Any additional information required to reanalyze the data reported in this paper is available from the lead contact upon request.

## Notes

### Competing Interest Statement

The authors have declared no competing interest.

### Funding

- National Institute of Health: 2T32EY007143, R01EY036173, R01EY031685, R01EY030872
- BrightFocus Foundation: G2019300
- Maryland Technology Development Corporation: 2022-MSCRFD-5895

## References

1. Agarwal, D., et al. Human retinal ganglion cell neurons generated by synchronous BMP inhibition and transcription factor mediated reprogramming. NPJ Regen Med. 2023;8(1):55. doi: 10.1038/s41536-023-00327-x.

2. Aijaz, S., et al. Expression analysis of SIX3 and SIX6 in human tissues reveals differences in expression and a novel correlation between the expression of SIX3 and the genes encoding isocitrate dehyhrogenase and cadherin 18. Genomics. 2005;86(1):86–99. doi: 10.1016/j.ygeno.2005.03.002.

3. Aldiri, I., et al. The Dynamic Epigenetic Landscape of the Retina During Development, Reprogramming, and Tumorigenesis. Neuron. 2017;94(3):550–68 e10. doi: 10.1016/j.neuron.2017.04.022.

4. Alexiades, M.R., Cepko, C. Quantitative analysis of proliferation and cell cycle length during development of the rat retina. Dev Dyn. 1996;205(3):293–307. doi: 10.1002/(SICI)1097-0177(199603)205:3<293::AID-AJA9>3.0.CO;2-D.

5. Berger, E., et al. Millifluidic culture improves human midbrain organoid vitality and differentiation. Lab Chip. 2018;18(20):3172–83. doi: 10.1039/c8lc00206a.

6. Bozanic, D., et al. Role of apoptosis and mitosis during human eye development. Eur J Cell Biol. 2003;82(8):421–9. doi: 10.1078/0171-9335-00328.

7. Breiman, L. Random forests. Mach Learn. 2001;45(1):5–32. doi: Doi 10.1023/A:1010933404324.

8. Brown, N.L., et al. Molecular characterization and mapping of ATOH7, a human atonal homolog with a predicted role in retinal ganglion cell development. Mamm Genome. 2002;13(2):95–101. doi: 10.1007/s00335-001-2101-3.

9. Brzezinski, J.A.t., et al. Math5 defines the ganglion cell competence state in a subpopulation of retinal progenitor cells exiting the cell cycle. Dev Biol. 2012;365(2):395–413. doi: 10.1016/j.ydbio.2012.03.006.

10. Buss, R.R., et al. Adaptive roles of programmed cell death during nervous system development. Annu Rev Neurosci. 2006;29:1–35. doi: 10.1146/annurev.neuro.29.051605.112800.

11. Candal, E., et al. Patterns of cell proliferation and cell death in the developing retina and optic tectum of the brown trout. Brain Res Dev Brain Res. 2005;154(1):101–19. doi: 10.1016/j.devbrainres.2004.10.008.

12. Capowski, E.E., et al. Reproducibility and staging of 3D human retinal organoids across multiple pluripotent stem cell lines. Development. 2019;146(1). doi: 10.1242/dev.171686.

13. Carpenter, P., et al. Role of target tissue in regulating the development of retinal ganglion cells in the albino rat: effects of kainate lesions in the superior colliculus. J Comp Neurol. 1986;251(2):240–59. doi: 10.1002/cne.902510208.

14. Cecconi, F., et al. Apaf1 (CED-4 homolog) regulates programmed cell death in mammalian development. Cell. 1998;94(6):727–37. doi: 10.1016/s0092-8674(00)81732-8.

15. Chavarria, T., et al. Differential, age-dependent MEK-ERK and PI3K-Akt activation by insulin acting as a survival factor during embryonic retinal development. Dev Neurobiol. 2007;67(13):1777–88. doi: 10.1002/dneu.20554.

16. Clark, B.S., et al. Single-Cell RNA-Seq Analysis of Retinal Development Identifies NFI Factors as Regulating Mitotic Exit and Late-Born Cell Specification. Neuron. 2019;102(6):1111–26 e5. doi: 10.1016/j.neuron.2019.04.010.

17. Cook, B., et al. Developmental neuronal death is not a universal phenomenon among cell types in the chick embryo retina. J Comp Neurol. 1998;396(1):12–9. doi: 10.1002/(sici)1096-9861(19980622)396:1<12::aid-cne2>3.0.co;2-l.

18. Cooper, E.R. The development of the human lateral geniculate body. Brain. 1945;68(3):222–39. doi: 10.1093/brain/68.3.222.

19. Costigan, A., et al. Discriminating Between Apoptosis, Necrosis, Necroptosis, and Ferroptosis by Microscopy and Flow Cytometry. Curr Protoc. 2023;3(12):e951. doi: 10.1002/cpz1.951.

20. Cowan, C.S., et al. Cell Types of the Human Retina and Its Organoids at Single-Cell Resolution. Cell. 2020;182(6):1623–40 e34. doi: 10.1016/j.cell.2020.08.013.

21. Crowley, L.C., et al. Measuring Cell Death by Propidium Iodide Uptake and Flow Cytometry. Cold Spring Harb Protoc. 2016;2016(7). doi: 10.1101/pdb.prot087163.

22. Diaz, B., et al. In vivo regulation of cell death by embryonic (pro)insulin and the insulin receptor during early retinal neurogenesis. Development. 2000;127(8):1641–9. doi: 10.1242/dev.127.8.1641.

23. Drabbe, E., et al. Retinal organoid chip: engineering a physiomimetic oxygen gradient for optimizing long term culture of human retinal organoids. Lab Chip. 2025;25(7):1626–36. doi: 10.1039/d4lc00771a.

24. Eldred, K.C., et al. Thyroid hormone signaling specifies cone subtypes in human retinal organoids. Science. 2018;362(6411). doi: 10.1126/science.aau6348.

25. Faure, L., et al. scFates: a scalable python package for advanced pseudotime and bifurcation analysis from single-cell data. Bioinformatics. 2023;39(1). doi: 10.1093/bioinformatics/btac746.

26. Fligor, C.M., et al. Three-Dimensional Retinal Organoids Facilitate the Investigation of Retinal Ganglion Cell Development, Organization and Neurite Outgrowth from Human Pluripotent Stem Cells. Sci Rep. 2018;8(1):14520. doi: 10.1038/s41598-018-32871-8.

27. Fligor, C.M., et al. Extension of retinofugal projections in an assembled model of human pluripotent stem cell-derived organoids. Stem Cell Reports. 2021;16(9):2228–41. doi: 10.1016/j.stemcr.2021.05.009.

28. Frade, J.M. Unscheduled re-entry into the cell cycle induced by NGF precedes cell death in nascent retinal neurones. J Cell Sci. 2000;113 (Pt 7):1139–48. doi: 10.1242/jcs.113.7.1139.

29. Frade, J.M., et al. Control of early cell death by BDNF in the chick retina. Development. 1997;124(17):3313–20. doi: 10.1242/dev.124.17.3313.

30. Frade, J.M., et al. Induction of cell death by endogenous nerve growth factor through its p75 receptor. Nature. 1996;383(6596):166–8. doi: 10.1038/383166a0.

31. Francisco-Morcillo, J., et al. Ontogenetic cell death and phagocytosis in the visual system of vertebrates. Dev Dyn. 2014;243(10):1203–25. doi: 10.1002/dvdy.24174.

32. Gaze, R.M., Grant, P. Spatio-temporal patterns of retinal ganglion cell death during Xenopus development. J Comp Neurol. 1992;315(3):264–74. doi: 10.1002/cne.903150303.

33. Georges, P., et al. Apoptosis during development of the human retina: relationship to foveal development and retinal synaptogenesis. J Comp Neurol. 1999;413(2):198–208.

34. Giandomenico, S.L., et al. Cerebral organoids at the air-liquid interface generate diverse nerve tracts with functional output. Nat Neurosci. 2019;22(4):669–79. doi: 10.1038/s41593-019-0350-2.

35. Gilbert, M.S. The early development of the human diencephalon. Journal of Comparative Neurology. 1935;62(1):81–115. doi: DOI 10.1002/cne.900620105.

36. Glucksmann, A. Development and Differentiation of the Tadpole Eye. Br J Ophthalmol. 1940;24(4):153–78. doi: 10.1136/bjo.24.4.153.

37. Gong, J., et al. A controllable perfusion microfluidic chip for facilitating the development of retinal ganglion cells in human retinal organoids. Lab Chip. 2023;23(17):3820–36. doi: 10.1039/d3lc00054k.

38. Guy, B., et al. Human neural organoids: Models for developmental neurobiology and disease. Dev Biol. 2021;478:102–21. doi: 10.1016/j.ydbio.2021.06.012.

39. Hahn, P., et al. Proapoptotic bcl-2 family members, Bax and Bak, are essential for developmental photoreceptor apoptosis. Invest Ophthalmol Vis Sci. 2003;44(8):3598–605. doi: 10.1167/iovs.02-1113.

40. Haskins, J.S., et al. Evaluating the Genotoxic and Cytotoxic Effects of Thymidine Analogs, 5-Ethynyl-2’-Deoxyuridine and 5-Bromo-2’-Deoxyurdine to Mammalian Cells. Int J Mol Sci. 2020;21(18). doi: 10.3390/ijms21186631.

41. Hensey, C., Gautier, J. Programmed cell death during Xenopus development: a spatio-temporal analysis. Dev Biol. 1998;203(1):36–48. doi: 10.1006/dbio.1998.9028.

42. Hevner, R.F. Development of connections in the human visual system during fetal mid-gestation: a DiI-tracing study. J Neuropathol Exp Neurol. 2000;59(5):385–92. doi: 10.1093/jnen/59.5.385.

43. Hoshino, A., et al. Molecular Anatomy of the Developing Human Retina. Dev Cell. 2017;43(6):763–79 e4. doi: 10.1016/j.devcel.2017.10.029.

44. Hughes, W.F., McLoon, S.C. Ganglion cell death during normal retinal development in the chick: comparisons with cell death induced by early target field destruction. Exp Neurol. 1979;66(3):587–601. doi: 10.1016/0014-4886(79)90204-8.

45. Hussey, K.A., et al. Patterning and Development of Photoreceptors in the Human Retina. Front Cell Dev Biol. 2022;10:878350. doi: 10.3389/fcell.2022.878350.

46. Inagaki, S., et al. Establishment of vascularized human retinal organoids from induced pluripotent stem cells. Stem Cells. 2025;43(3). doi: 10.1093/stmcls/sxae093.

47. Johnson, J.A.I., et al. Inferring cellular and molecular processes in single-cell data with non-negative matrix factorization using Python, R and GenePattern Notebook implementations of CoGAPS. Nat Protoc. 2023;18(12):3690–731. doi: 10.1038/s41596-023-00892-x.

48. Joshi, P., et al. Modeling the function of BAX and BAK in early human brain development using iPSC-derived systems. Cell Death Dis. 2020;11(9):808. doi: 10.1038/s41419-020-03002-x.

49. Kanda, T., et al. Histone-GFP fusion protein enables sensitive analysis of chromosome dynamics in living mammalian cells. Curr Biol. 1998;8(7):377–85. doi: 10.1016/s0960-9822(98)70156-3.

50. Kaur, C., et al. Hypoxia-ischemia and retinal ganglion cell damage. Clin Ophthalmol. 2008;2(4):879–89. doi: 10.2147/opth.s3361.

51. Kay, J.N., et al. Staggered cell-intrinsic timing of ath5 expression underlies the wave of ganglion cell neurogenesis in the zebrafish retina. Development. 2005;132(11):2573–85. doi: 10.1242/dev.01831.

52. Ke, F.F.S., et al. Embryogenesis and Adult Life in the Absence of Intrinsic Apoptosis Effectors BAX, BAK, and BOK. Cell. 2018;173(5):1217–30 e17. doi: 10.1016/j.cell.2018.04.036.

53. Khan, A.A., et al. Development of human lateral geniculate nucleus: an electron microscopic study. Int J Dev Neurosci. 1994;12(7):661–72. doi: 10.1016/0736-5748(94)90018-3.

54. Kretz, A., et al. Regulation of GDNF and its receptor components GFR-alpha1, -alpha2 and Ret during development and in the mature retino-collicular pathway. Brain Res. 2006;1090(1):1–14. doi: 10.1016/j.brainres.2006.01.131.

55. Kriukov, E., et al. Unraveling the developmental heterogeneity within the human retina to reconstruct the continuity of retinal ganglion cell maturation and stage-specific intrinsic and extrinsic factors. bioRxiv. 2024. doi: 10.1101/2024.10.16.618776.

56. Laha, B., et al. Regenerating optic pathways from the eye to the brain. Science. 2017;356(6342):1031-4. doi: 10.1126/science.aal5060.

57. Lancaster, M.A., et al. Cerebral organoids model human brain development and microcephaly. Nature. 2013;501(7467):373-9. doi: 10.1038/nature12517.

58. Langer, K.B., et al. Retinal Ganglion Cell Diversity and Subtype Specification from Human Pluripotent Stem Cells. Stem Cell Reports. 2018;10(4):1282–93. doi: 10.1016/j.stemcr.2018.02.010.

59. Liu, S.B., et al. G-protein-coupled receptor 30 mediates rapid neuroprotective effects of estrogen via depression of NR2B-containing NMDA receptors. J Neurosci. 2012;32(14):4887–900. doi: 10.1523/JNEUROSCI.5828-11.2012.

60. Liu, Y.V., et al. Single-cell transcriptome analysis of xenotransplanted human retinal organoids defines two migratory cell populations of nonretinal origin. Stem Cell Reports. 2023;18(5):1138–54. doi: 10.1016/j.stemcr.2023.04.004.

61. Lom, B., Cohen-Cory, S. Brain-derived neurotrophic factor differentially regulates retinal ganglion cell dendritic and axonal arborization in vivo. J Neurosci. 1999;19(22):9928–38. doi: 10.1523/JNEUROSCI.19-22-09928.1999.

62. Lu, Y., et al. Single-Cell Analysis of Human Retina Identifies Evolutionarily Conserved and Species-Specific Mechanisms Controlling Development. Dev Cell. 2020;53(4):473–91 e9. doi: 10.1016/j.devcel.2020.04.009.

63. Luo, Z., et al. Islet1 and Brn3 Expression Pattern Study in Human Retina and hiPSC-Derived Retinal Organoid. Stem Cells Int. 2019;2019:8786396. doi: 10.1155/2019/8786396.

64. Mayordomo, R., et al. Generation of retinal ganglion cells is modulated by caspase-dependent programmed cell death. Eur J Neurosci. 2003;18(7):1744–50. doi: 10.1046/j.1460-9568.2003.02891.x.

65. McLaughlin, T., et al. Retinotopic map refinement requires spontaneous retinal waves during a brief critical period of development. Neuron. 2003;40(6):1147–60. doi: 10.1016/s0896-6273(03)00790-6.

66. McNerney, C., et al. DIO3 coordinates photoreceptor development timing and fate stability in human retinal organoids. bioRxiv. 2025:2025.03. 20.644422.

67. Meyer, J.S., et al. Optic vesicle-like structures derived from human pluripotent stem cells facilitate a customized approach to retinal disease treatment. Stem Cells. 2011;29(8):1206–18. doi: 10.1002/stem.674.

68. Mu, X., et al. Ganglion cells are required for normal progenitor-cell proliferation but not cell-fate determination or patterning in the developing mouse retina. Curr Biol. 2005;15(6):525–30. doi: 10.1016/j.cub.2005.01.043.

69. Nakano, T., et al. Self-formation of optic cups and storable stratified neural retina from human ESCs. Cell Stem Cell. 2012;10(6):771–85. doi: 10.1016/j.stem.2012.05.009.

70. Nguyen-Ba-Charvet, K.T., Rebsam, A. Neurogenesis and Specification of Retinal Ganglion Cells. Int J Mol Sci. 2020;21(2). doi: 10.3390/ijms21020451.

71. Oppenheim, R. Neuronal cell death and some related regressive phenomena during neurogenesis: a selective historical review and progress report. Studies in developmental neurobiology. 1981:74–133.

72. Osborne, N.N., et al. Retinal ischemia: mechanisms of damage and potential therapeutic strategies. Prog Retin Eye Res. 2004;23(1):91–147. doi: 10.1016/j.preteyeres.2003.12.001.

73. Ovando-Roche, P., et al. Use of bioreactors for culturing human retinal organoids improves photoreceptor yields. Stem Cell Res Ther. 2018;9(1):156. doi: 10.1186/s13287-018-0907-0.

74. Pena-Blanco, A., Garcia-Saez, A.J. Bax, Bak and beyond - mitochondrial performance in apoptosis. FEBS J. 2018;285(3):416–31. doi: 10.1111/febs.14186.

75. Pequignot, M.O., et al. Major role of BAX in apoptosis during retinal development and in establishment of a functional postnatal retina. Dev Dyn. 2003;228(2):231–8. doi: 10.1002/dvdy.10376.

76. Perry, V.H., Cowey, A. A sensitive period for ganglion cell degeneration and the formation of aberrant retino-fugal connections following tectal lesions in rats. Neuroscience. 1982;7(3):583–94. doi: 10.1016/0306-4522(82)90065-3.

77. Porter, A.G., Janicke, R.U. Emerging roles of caspase-3 in apoptosis. Cell Death Differ. 1999;6(2):99–104. doi: 10.1038/sj.cdd.4400476.

78. Provis, J.M. Patterns of cell death in the ganglion cell layer of the human fetal retina. J Comp Neurol. 1987;259(2):237–46. doi: 10.1002/cne.902590205.

79. Provis, J.M., et al. Ganglion cell topography in human fetal retinae. Invest Ophthalmol Vis Sci. 1983;24(9):1316–20.

80. Provis, J.M., et al. Development of the human retina: patterns of cell distribution and redistribution in the ganglion cell layer. J Comp Neurol. 1985a;233(4):429–51. doi: 10.1002/cne.902330403.

81. Provis, J.M., et al. Human fetal optic nerve: overproduction and elimination of retinal axons during development. J Comp Neurol. 1985b;238(1):92–100. doi: 10.1002/cne.902380108.

82. Provis, J.M., Vandriel, D. Retinal Development in Humans - the Roles of Differential Growth-Rates, Cell-Migration and Naturally-Occurring Cell-Death. Aust Nz J Ophthalmol. 1985;13(2):125–33. doi: DOI 10.1111/j.1442-9071.1985.tb00413.x.

83. Qian, X., et al. Sliced Human Cortical Organoids for Modeling Distinct Cortical Layer Formation. Cell Stem Cell. 2020;26(5):766–81 e9. doi: 10.1016/j.stem.2020.02.002.

84. Quinn, P.M.J., Wijnholds, J. Retinogenesis of the Human Fetal Retina: An Apical Polarity Perspective. Genes (Basel). 2019;10(12). doi: 10.3390/genes10120987.

85. Rakic, P., Riley, K.P. Overproduction and elimination of retinal axons in the fetal rhesus monkey. Science. 1983;219(4591):1441-4. doi: 10.1126/science.6828871.

86. Rieger, A.M., et al. Modified annexin V/propidium iodide apoptosis assay for accurate assessment of cell death. J Vis Exp. 2011(50). doi: 10.3791/2597.

87. Rodieck, R.W., Marshak, D.W. Spatial density and distribution of choline acetyltransferase immunoreactive cells in human, macaque, and baboon retinas. J Comp Neurol. 1992;321(1):46–64. doi: 10.1002/cne.903210106.

88. Rodriguez, A.R., et al. The RNA binding protein RBPMS is a selective marker of ganglion cells in the mammalian retina. J Comp Neurol. 2014;522(6):1411–43. doi: 10.1002/cne.23521.

89. Shiau, F., et al. A single-cell guide to retinal development: Cell fate decisions of multipotent retinal progenitors in scRNA-seq. Dev Biol. 2021;478:41–58. doi: 10.1016/j.ydbio.2021.06.005.

90. Sluch, V.M., et al. Enhanced Stem Cell Differentiation and Immunopurification of Genome Engineered Human Retinal Ganglion Cells. Stem Cells Transl Med. 2017;6(11):1972–86. doi: 10.1002/sctm.17-0059.

91. Soto, I., et al. Retinal ganglion cells downregulate gene expression and lose their axons within the optic nerve head in a mouse glaucoma model. J Neurosci. 2008;28(2):548–61. doi: 10.1523/JNEUROSCI.3714-07.2008.

92. Soucy, J.R., et al. Retinal ganglion cell repopulation for vision restoration in optic neuropathy: a roadmap from the RReSTORe Consortium. Mol Neurodegener. 2023;18(1):64. doi: 10.1186/s13024-023-00655-y.

93. Stein-O’Brien, G.L., et al. Decomposing Cell Identity for Transfer Learning across Cellular Measurements, Platforms, Tissues, and Species. Cell Syst. 2019;8(5):395–411 e8. doi: 10.1016/j.cels.2019.04.004.

94. Surgucheva, I., et al. Gamma-synuclein as a marker of retinal ganglion cells. Mol Vis. 2008;14:1540–8.

95. Surguchov, A., et al. Synucleins in ocular tissues. J Neurosci Res. 2001;65(1):68–77. doi: 10.1002/jnr.1129.

96. Tayler, K.K., et al. Characterization of NMDAR-Independent Learning in the Hippocampus. Front Behav Neurosci. 2011;5:28. doi: 10.3389/fnbeh.2011.00028.

97. Tian, N., Copenhagen, D.R. Visual stimulation is required for refinement of ON and OFF pathways in postnatal retina. Neuron. 2003;39(1):85–96. doi: 10.1016/s0896-6273(03)00389-1.

98. Toy, J., Sundin, O.H. Expression of the optx2 homeobox gene during mouse development. Mech Dev. 1999;83(1-2):183–6. doi: 10.1016/s0925-4773(99)00049-0.

99. Toy, J., et al. The optx2 homeobox gene is expressed in early precursors of the eye and activates retina-specific genes. Proc Natl Acad Sci U S A. 1998;95(18):10643–8. doi: 10.1073/pnas.95.18.10643.

100. Trimarchi, J.M., et al. Molecular heterogeneity of developing retinal ganglion and amacrine cells revealed through single cell gene expression profiling. J Comp Neurol. 2007;502(6):1047–65. doi: 10.1002/cne.21368.

101. Vanselow, J., et al. Target dependence of chick retinal ganglion cells during embryogenesis: cell survival and dendritic development. J Comp Neurol. 1990;295(2):235–47. doi: 10.1002/cne.902950207.

102. Volkner, M., et al. Mouse Retinal Organoid Growth and Maintenance in Longer-Term Culture. Front Cell Dev Biol. 2021;9:645704. doi: 10.3389/fcell.2021.645704.

103. Wahle, P., et al. Multimodal spatiotemporal phenotyping of human retinal organoid development. Nat Biotechnol. 2023;41(12):1765–75. doi: 10.1038/s41587-023-01747-2.

104. Wahlin, K.J., et al. CRISPR Generated SIX6 and POU4F2 Reporters Allow Identification of Brain and Optic Transcriptional Differences in Human PSC-Derived Organoids. Front Cell Dev Biol. 2021;9:764725. doi: 10.3389/fcell.2021.764725.

105. Wahlin, K.J., et al. Photoreceptor Outer Segment-like Structures in Long-Term 3D Retinas from Human Pluripotent Stem Cells. Sci Rep. 2017;7(1):766. doi: 10.1038/s41598-017-00774-9.

106. Wang, Q., et al. Ipsilateral and Contralateral Retinal Ganglion Cells Express Distinct Genes during Decussation at the Optic Chiasm. eNeuro. 2016;3(6). doi: 10.1523/ENEURO.0169-16.2016.

107. Wang, Y., et al. Retinal ganglion cell-derived sonic hedgehog locally controls proliferation and the timing of RGC development in the embryonic mouse retina. Development. 2005;132(22):5103–13. doi: 10.1242/dev.02096.

108. Whitney, I.E., et al. Vision-Dependent and -Independent Molecular Maturation of Mouse Retinal Ganglion Cells. Neuroscience. 2023;508:153–73. doi: 10.1016/j.neuroscience.2022.07.013.

109. Wiese, C.B., et al. A Uchl1-Histone2BmCherry:GFP-gpi BAC transgene for imaging neuronal progenitors. Genesis. 2013;51(12):852–61. doi: 10.1002/dvg.22716.

110. Wolf, F.A., et al. SCANPY: large-scale single-cell gene expression data analysis. Genome Biol. 2018;19(1):15. doi: 10.1186/s13059-017-1382-0.

111. Xue, Y., et al. The Prospects for Retinal Organoids in Treatment of Retinal Diseases. Asia Pac J Ophthalmol (Phila). 2022;11(4):314–27. doi: 10.1097/APO.0000000000000538.

112. Yamaguchi, Y., Miura, M. Programmed cell death in neurodevelopment. Dev Cell. 2015;32(4):478–90. doi: 10.1016/j.devcel.2015.01.019.

113. Yan, W., et al. Cell Atlas of The Human Fovea and Peripheral Retina. Sci Rep. 2020;10(1):9802. doi: 10.1038/s41598-020-66092-9.

114. Young, R.W. Cell death during differentiation of the retina in the mouse. J Comp Neurol. 1984;229(3):362–73. doi: 10.1002/cne.902290307.

115. Zhang, K.Y., Johnson, T.V. The internal limiting membrane: Roles in retinal development and implications for emerging ocular therapies. Exp Eye Res. 2021;206:108545. doi: 10.1016/j.exer.2021.108545.

116. Zhong, X., et al. Generation of three-dimensional retinal tissue with functional photoreceptors from human iPSCs. Nat Commun. 2014;5:4047. doi: 10.1038/ncomms5047.

117. Zuo, Z., et al. Single cell dual-omic atlas of the human developing retina. Nat Commun. 2024;15(1):6792. doi: 10.1038/s41467-024-50853-5.

